# The chromatin remodeler CHD3 is highly expressed in mature neurons and regulates genes involved in synaptic development and function

**DOI:** 10.1101/2024.04.29.591720

**Authors:** Joery den Hoed, Maggie M. K. Wong, Willemijn J. J. Claassen, Lucía de Hoyos, Lukas Lütje, Michael Heide, Wieland B. Huttner, Simon E. Fisher

**Affiliations:** Language and Genetics Department, Max Planck Institute for Psycholinguistics, Nijmegen, the Netherlands; Max Planck Institute of Molecular Cell Biology and Genetics, Dresden, Germany; Donders Institute for Brain, Cognition and Behaviour, Radboud University, Nijmegen, The Netherlands

**Author notes:** To whom correspondence should be addressed: Prof. Dr. S.E. Fisher. German Primate Center, Leibniz Institute for Primate Research, Göttingen, Germany (Michael Heide).

## Abstract

Changes in the dynamics of chromatin state that control spatiotemporal gene expression patterns are crucial during brain development. CHD3 is a chromatin remodeler that is highly expressed during neurogenesis and that functions as a core member of the NuRD complex, a large multiprotein complex mediating chromatin state. Genetic disruptions in *CHD3* have been implicated in a neurodevelopmental disorder characterized by intellectual disability, macrocephaly and severe speech deficits. To study the roles of CHD3 during early human brain development, we generated induced pluripotent stem cells with heterozygous and homozygous loss-of-function mutations, differentiated them into unguided neural organoids and cortical neurons, and analyzed these by immunohistochemistry, bulk RNA-, single-cell RNA-, and ChIP-sequencing. Loss of *CHD3* expression had no detectable effects on early neuroepithelium formation and organoid growth, nor did it significantly affect cell type composition or neuronal differentiation speed. Instead, upon loss of *CHD3*, we observed dysregulation of genes related to axon guidance and synapse development across all datasets, identifying a novel role for the protein as a regulator that facilitates neurogenesis, in particular neuronal maturation. Our results based on genetically engineered knockout organoids pave the way for future studies modeling the neurobiological pathways affected in CHD3-related disorder.

## Introduction

The Chromodomain Helicase DNA-binding protein (CHD) family is a group of chromatin remodelers that have emerged as important regulators of brain development. From its nine members, divided in three subfamilies, seven have been implicated in neurodevelopmental disorders^1–7^.

One of the most recently described of these disorders is a broad neurodevelopmental syndrome associated with heterozygous *de novo* missense variants in *CHD3*, mostly clustering in the ATPase-Helicase domain crucial for the protein’s chromatin remodeling function (Snijders Blok-Campeau syndrome, SNIBCPS, MIM 618205)^3^. Since then, the cohort of known and described cases has been further extended, confirming the overrepresentation of missense variants located in the ATPase-Helicase domain^8,9^. Functional analyses demonstrated that at least a subset of these missense variants affect either the ATPase activity of CHD3, its nucleosome shuffling capacity, or both^3^. Moreover, heterozygous protein-truncating variants of CHD3 were subsequently shown to be associated with neurodevelopmental phenotypes similar to the missense variants, albeit often with variable expressivity^10^. Despite its association with neurodevelopmental disorder, the functions of CHD3 during human brain development and the effects of etiological variants in this context remain largely undetermined.

CHD3, CHD4 and CHD5 belong to a single subfamily of the CHD family, sharing all their known functional domains^11^, and serve as one of the core subunits of a large protein complex with both histone deacetylase and chromatin remodeling activities, called the NuRD (**Nu**cleosome **R**emodeling and **D**eacetylase) complex^12^. From this subfamily, CHD4 has been most intensively studied in the context of brain development, in particular in the mouse cerebellum^13^. Here, the Chd4-associated NuRD complex drives synapse formation of granule neurons and Purkinje cells, and consequently, *Chd4* disruption results in a reduced number of synapses in these cells^14^. While the Chd4 protein was found to negatively regulate about 200 direct target genes in the mouse cerebellum^14^, it also occupies regulatory regions of actively transcribed genes, where it depositions the histone variant H2A.Z over H2A. This way, CHD4 indirectly controls the expression of immediate early genes that are involved in the pruning of granule neuron dendrites^15^, demonstrating that the protein is essential for multiple distinct processes linked to neuronal connectivity in the cerebellum.

A study focusing on this subfamily of Chd proteins in mouse neocortical development, demonstrated that Chd3, Chd4 and Chd4 are mutually exclusive in the NuRD complex, and have non-redundant roles^12^. Distinct CHD4-NuRD and CHD3-NuRD complexes with non-overlapping functions have also been confirmed in human cell lines^16^. In the mouse neocortex, Chd4 seems to be most crucial at early stages of cortical development, promoting the proliferation of neural progenitor cells (NPCs) by activating the expression of *Pax6*, *Sox2* and *Tbr2/Eomes*. When Chd3 and Chd5 replace Chd4, the function of the NuRD complex changes and neuronal differentiation is induced. Chd3 facilitates radial migration coupled with late cortical layer specification by repressing *Pax6*, *Sox2* and *Tbr2/Eomes*. On the other hand, Chd5 seems crucial for earlier stages of neuronal migration, activating regulators of neuronal differentiation such as *Dcx* and *RhoA*, and *Chd5* knockdown causes the formation of an ectopic layer of mature neurons in the ventricular zone^12^. A separate study focusing only on *Chd5* observed slightly different effects upon knockdown of this gene in the mouse cortex, resulting in an accumulation of NPCs that fail to exit the cell cycle and continue to express NPC marker genes, fully disrupting normal neuronal differentiation^17^.

Although work with mouse models has shown the importance of the NuRD complex-associated CHD proteins in cerebellar and cortical development^13,18^, in particular in controlling the transition towards neuronal differentiation and migration, their roles in the ontogeny of the human brain and their links with human disease are still poorly understood. In the current study, we aimed to investigate this question, by using a combination of CRISPR-Cas9 gene-edited stem cell lines with *CHD3* knockout mutations and the generation of stem cell-derived neural organoids and forebrain neurons to model early stages of human brain development. Overall, we did not observe gross differences in neural organoid growth and ventricle-like structures, nor disturbance in formation of MAP2-positive forebrain neurons, when *CHD3* was disrupted. Following up with single-cell transcriptomic analyses, we show that *CHD3* is not an essential factor for inducing and promoting neuronal differentiation, as all cell types detected in the neural organoid model, including post-mitotic neurons, were still represented in similar proportions in complete *CHD3* knockouts as compared to wild-type controls. Against this background, both differential gene expression analyses and chromatin immunoprecipitation (ChIP) sequencing data converge on a previously unknown role for CHD3 more specifically in mature neurons, as a regulator of genes associated with synapse development and function, and axon guidance and growth.

## Results

### CRISPR-induced frameshift variants in exon 3 disrupt *CHD3* expression in induced pluripotent stem cells and unguided neural organoids

We sought to investigate the functions of CHD3 during human brain development by employing a stem cell-based organoid model system. In analyses of human developmental transcriptomic data of sub-dissected cortical regions from BrainSpan^19^, we found *CHD3* to have peak expression levels throughout all cortical regions at 16-19 weeks post-conception (Figure S1). Therefore, we decided to perform our investigations of CHD3 function using a widely used and well-established brain organoid model that recapitulates aspects of *in vivo* cortical development: cerebral organoids^20,21^. Following the guidelines in a recent proposal on the nomenclature consensus for organoids modeling the nervous system^22^, we refer to the model used in this study as “unguided neural organoids”.

We employed CRISPR-Cas9 gene editing to generate heterozygous, compound heterozygous and homozygous *CHD3* loss-of-function frameshift variants in a commercially available human induced pluripotent stem cell line from a healthy donor (BIONi010-A; 010A^P^; iPSC)^23^. We generated three clonal cell lines for each *CHD3* genotype (010A^WT/KO^ and 010A^KO/KO^), as well as clonal cell lines that underwent the gene editing procedure without the introduction of mutations at the target site (010A^WT/WT^; Figure 1A). This involved use of a guide-RNA that induced a double-strand DNA break in exon 3 of *CHD3*, thereby targeting all three of the major isoforms (Figure 1B). Nonhomologous end joining caused either a 1-bp deletion (in cell lines 010A^WT/KO^ C1, 010A^KO/KO^ C1, 010A^KO/KO^ C2; c.298delG, p.G100Vfs*40; NM_001005273.2), or a 1-bp insertion (in cell line 010A^WT/KO^ C2; c.298insA, p.G100Efs*54 and 010A^WT/KO^ C3; c.298insT, p.G100Vfs*54) at the target site (Figure 1C), in each case resulting in the formation of a premature stop-codon downstream. Cell line 010A^KO/KO^ C3 was compound heterozygous, with a c.298delG (p.G100Vfs*40) and a c.298insG (p.R101Sfs*53) mutation on either allele (Figure 1C).

**Figure 1.**
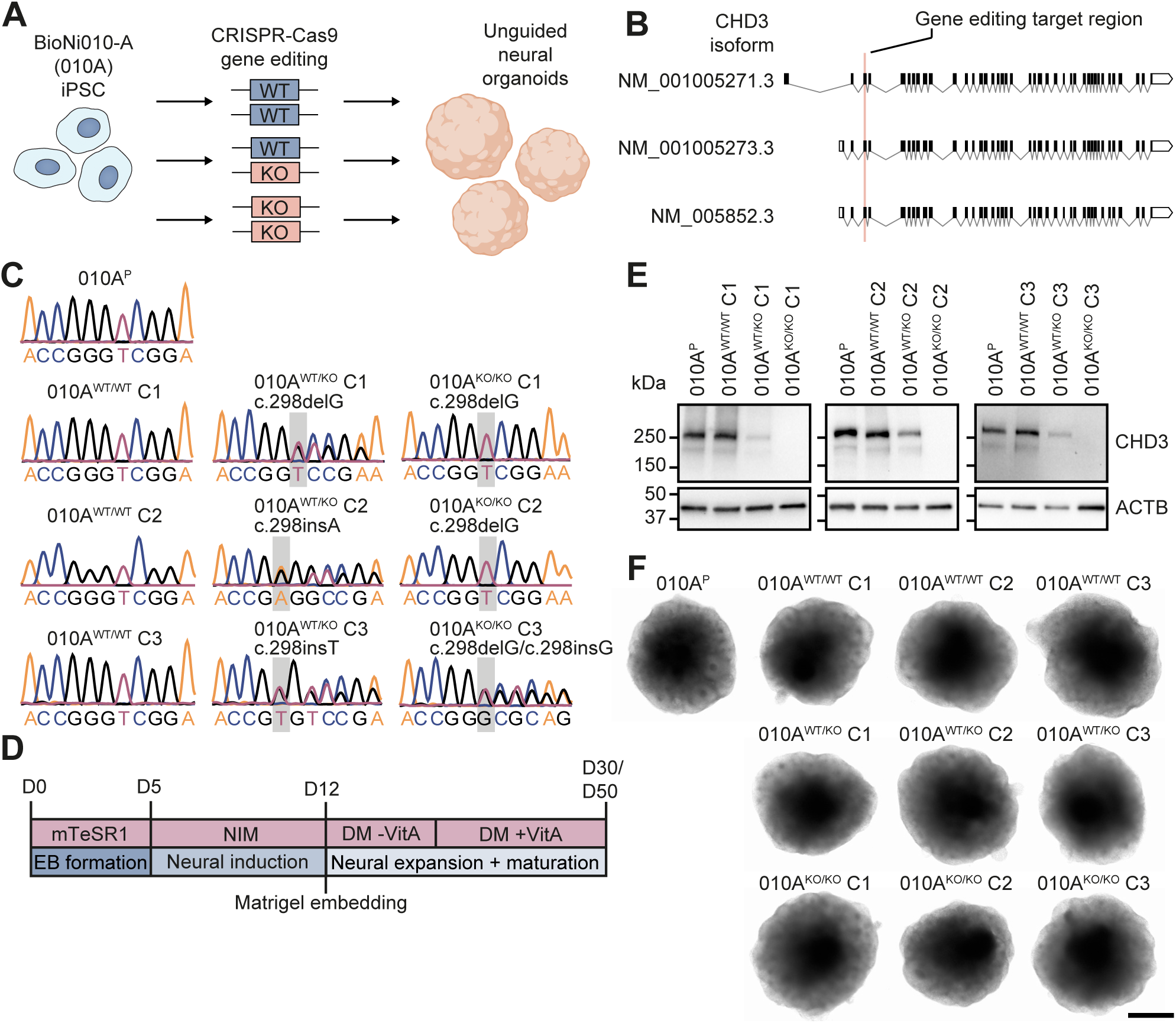
CRISPR-induced frameshift variants in exon 3 disrupt *CHD3* expression in unguided neural organoids. **A)** Overview of the gene-editing approach used in this study, creating control cell lines that remained unedited after the CRISPR-Cas9 procedure and cell lines carrying a heterozygous, homozygous or compound heterozygous *CHD3* frameshift variant. **B**) Exon-intron schematics of the three major isoforms of *CHD3*, generated with the ExInPlotter tool (see Code Availability). The coding DNA sequence is shown in black, the 5’ and 3’ UTRs in white, and the introns are depicted with gray lines. The exon that was targeted by the CRISPR-Cas9 guide-RNA is highlighted with a red bar. **C**) Sanger traces of the CRISPR-Cas9 target region in clonal cell lines selected for further characterization. The unedited clones (010A^WT/WT^) show no differences from the parental target region (010A^P^), while the 010A^WT/KO^ and 010A^KO/KO^ clones carry a 1-bp indel, either in heterozygous state or on both alleles respectively. **D**) The unguided neural organoid culture protocol^20,21^, described in detail in the Methods section. **E**) Immunoblot of whole-cell lysates of three day-30 neural organoids pooled together expressing CHD3 protein. Expected molecular weight is ∼226 kDa. The blot was probed for ACTB to ensure equal protein loading. **F**) Representative bright-field images of day-15 unguided neural organoids for each cell line. Scale bar = 500 μm.

Selected clonal cell lines were tested for their chromosomal integrity using molecular karyotyping (Figure S2). All lines shared a 22q11.23 microduplication of approximately 1.3 Mb which was already present in the original cell line (010A^P^). Furthermore, 010A^WT/KO^ C2 and 010A^KO/KO^ C3 carried a 2 Mb 1q32.1 gain (Figure S2), a commonly found aberration in stem cell culturing^24^. Our gene-editing design predicted that there were nineteen potential CRISPR-Cas9 off-targets, of which only one was located in an exonic region (Table S1). We prioritized the exonic off-target, and randomly selected four intergenic predicted off-targets for screening with Sanger sequencing, establishing that they all remained unedited in the cell lines of the study (Figure S3). Next, we showed that CHD3 protein is already expressed at low levels in iPSCs and verified the knockout status of the 010A^WT/KO^ and 010A^KO/KO^ lines (Figure S4). All edited iPSC lines continued to express markers of pluripotency (OCT4, SOX2, SSEA-4 and TRA1-60) independent of the *CHD3* genotype, successfully confirming their stem cell state (Figure S5).

We went on to derive unguided neural organoids from these cell lines (010A^P^, 010A^WT/WT^ C1-3, 010A^WT/KO^ C1-3, and 010A^KO/KO^ C1-3) using methods previously described in the literature (Figure 1D)^20,21^. Analyzing day-30 organoids, we further confirmed reduced or absent CHD3 protein levels in 010A^WT/KO^ and 010A^KO/KO^ lines, showing a clear genotype-dependent dosage effect (Figure 1E). These findings suggest that the transcripts carrying the introduced *CHD3* mutations are subject to nonsense-mediated decay (as predicted based on the nature and location of the introduced variant) and show that CHD3 protein expression is disrupted.

### Unguided neural organoids lacking CHD3 expression have normal rates of growth

Next, we assessed the morphology and growth rates of neural organoids derived from cells lacking the expression of one or both *CHD3* alleles. Regardless of genotype, and with high consistency across independent clonal lines, all organoids had a similar appearance to each other during the first fifteen days of the protocol (Figure 1F and Figure S6). Measuring the surface area of the organoids, we did not find the *CHD3* genotype to influence the initial growth of the organoids during the first eleven days of organoid formation (Figure S6). These results show that CHD3 has limited impact on the earliest stages of neuroepithelium formation and initial expansion, consistent with our expectations that CHD3 plays a role later during neurodevelopment, based on the expression of the gene peaking at mid-gestation in the human developing cortex (Figure S1) and prior work in mouse models^12^.

### Unguided neural organoids lacking CHD3 expression contain both neural progenitors and mature neurons

Based on the hypothesis that CHD3 acts as a potential pro-neural factor^12^, we examined whether lower levels or a complete lack of CHD3 would affect the generation of mature neurons. We performed immunostainings on wild-type day-50 and day-57 organoids to characterize CHD3 expression in this cellular model and showed that the protein is particularly highly expressed in TUBB3-, CTIP2/BCL11A- and TBR1-positive mature neurons, while expression in TBR2/EOMES-positive intermediate progenitors and SOX2- and PAX6-positive radial glial cells is lower (Figure S7 and S8). In complete knockout organoids (010A^KO/KO^ C1-3), our immunostainings again confirmed a clear loss of CHD3 expression (Figure S9). However, organoids from all genotypes and consistently across independent clonal lines, still contained PAX6- and TBR1-positive cells organized in distinct subregions within the rosette-like structures, resembling the ventricular zone (PAX6-postive cells), the subventricular zone (PAX6-positive and TBR1-positive cells), and the early cortical plate (TBR1-positive cells; Figure S10). Although these data do not exclude the possibility of subtle differences in the numbers of NPCs and/or mature neurons and the organization of the rosettes between the genotypes, nor effects of decreased CHD3 expression on neuronal function, the staining results indicate that CHD3 is not essential for the differentiation of NPCs into CTIP2/BCL11A- and TBR1-positive neurons.

### Disruption of CHD3 does not affect organoid composition and timing of neuronal differentiation

To examine the cellular composition of the organoids, we performed single-cell RNA-seq for each cell line on day-56/57 organoids. We obtained data for 29,346 cells in total, of which 14,365 cells were from wild-type (representing three samples of 010A^P^, and one sample of 010A^WT/WT^ C1, 010A^WT/WT^ C2, and 010A^WT/WT^ C3), 5,997 cells from heterozygous CHD3 knockout (representing 010A^WT/KO^ C1, 010A^WT/KO^ C2, and 010A^WT/KO^ C3) and 8,984 cells from full CHD3 knockout organoids (representing 010A^KO/KO^ C1, 010A^KO/KO^ C2, and 010A^KO/KO^ C3) (Figure 2A; Figure S11A). We mapped our data onto a previously published dataset with 46,977 cells from 2-month old unguided neural organoids derived from seven different human stem cell lines cultured using the same protocol^25^, and transferred the cell type labels for annotation (Figure 2A-B). The identified cell types represented dorsal (20,927 cells), as well as ventral (7,333 cells) and non-telencephalic (1,086 cells) lineages (Figure 2C). Consistent with human developmental transcriptomic data (Figure S1) and prior studies in mice^12,19^, as well as our immunostainings in organoids (Figure S7 and S8), *CHD3* expression was higher in mature neurons [Cortical excitatory neurons (EN), medial/caudal ganglionic eminence (MGE/CGE) inhibitory neurons (IN), lateral ganglionic eminence (LGE) IN; Figure 2D] than in progenitor cell types [Cortical NPC, ganglionic eminence (GE) NPC, non-telencephalic (Non-tel) NPC; Figure 2D]. Moreover, as expected, *CHD3* transcript levels were decreased in cells with a heterozygous or complete CHD3 knockout mutation in a dosage-dependent manner in all dorsal and ventral cell types (Figure 2D).

**Figure 2.**
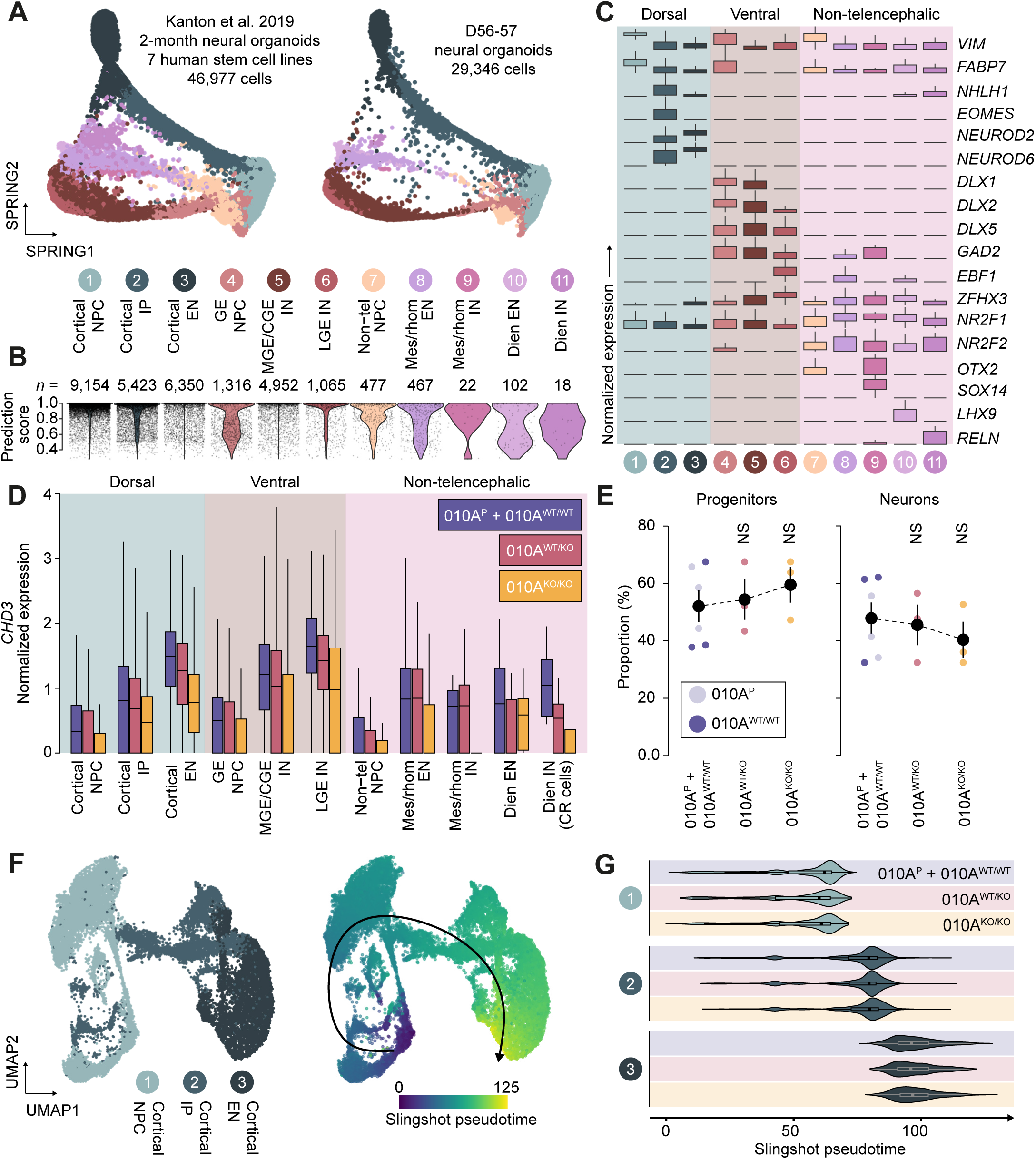
The effects of *CHD3* disruption on cell composition of unguided neural organoids. **A**) Left, the SPRING reconstruction of scRNA-sequencing data of 2-month old unguided neural organoids derived from seven human stem cell lines (46,977 cells) with clusters coloured by cell type, as described by Kanton et al. 2019, Ref. 24. Right, a scatter plot of 29,347 cells from day-56/57 unguided neural organoids derived from the gene-edited 010A cell lines. Cell types were annotated by querying the data on the dataset of Kanton et al. 2019, and cells were visualized in the same reduced dimensional space. CGE, caudal ganglionic eminence; EN, excitatory neuron; GE, ganglionic eminence; IN, inhibitory neuron; IP, intermediate progenitors; LGE, lateral ganglionic eminence; MGE, medial ganglionic eminence; NPC, neural progenitor cell. **B**) Violin plots of the prediction scores of the cell type annotations for each cell type after mapping and annotation of the query dataset. Total number of cells for each cell type in the query dataset are indicated. **C**) Box plots showing the normalized expression levels of a selection of marker genes for the different cell populations identified in the neural organoids. Dorsal cell types are shaded in blue, ventral cell types in red, and non-telencephalic cell types in pink. **D**) Boxplots of the normalized expression levels of *CHD3* for each cell type and across *CHD3* genotypes. Purple bars show the wild-type genotype, red the 010AWT/KO, and yellow the 010AKO/KO condition, shading as in (**C**). **E**) Relative cell type proportions across *CHD3* genotypes, with progenitors encompassing Cortical NPC, Cortical IP, GE NPC and non-tel NPC, and neurons including Cortical EN, MGE/CGE IN, LGE IN, Mes/rhom EN, Mes/rhom IN, Dien EN and Dien IN. The plot shows the mean proportion ± SEM. NS, not significant; one-way ANOVA and *post hoc* Dunnet test. Colouring as in (**D**). **F**) Left, a subset of the scRNA-sequencing data derived from the gene-edited 010A cell lines, only containing the dorsal cell types (Cortical NPC, Cortical IP, Cortical EN) and shown in a UMAP embedding. Right, the dorsal trajectory inferred by Slingshot and displayed on top of the subsetted data. **G**) Violin plots of Slingshot pseudotime values for each dorsal cell type across *CHD3* genotypes.

When we examined proportions of progenitor cells [Cortical NPC, Cortical intermediate progenitors (IP), GE NPC and Non-tel NPC; Figure 2] and neuronal cells [Cortical NE, MGE/CGE IN, LGE IN, mesencephalon/rhombencephalon (Mes/rhom) EN, Mes/rhom IN, diencephalon (Dien) EN and Dien IN; Figure 2] across the different cell lines, we did not detect significant differences related to *CHD3* genotype (Figure 2E; Figure S11B). After subsetting the dorsal cell types, the most represented lineage in our dataset (71.3% of cells; Figure 2F), and performing Slingshot pseudotime analysis^26^, we did not observe major shifts in the distribution of cells across the inferred dorsal pseudotime trajectory for the different genotypes (Figure 2G). These results suggest that CHD3 is not a crucial factor for neuronal differentiation in unguided neural organoids, nor does it seem to affect the timing and speed of differentiation.

### A selection of genes is differentially expressed in cortical excitatory neurons lacking *CHD3* expression

We then performed pseudobulk differential gene expression analysis on the dorsal (Cortical NPC, Cortical IP and Cortical EN) and ventral (GE NPC, MGE/CGE IN, LGE IN) cell types, comparing the complete *CHD3* knockout to the wild-type condition. Overall, we found only a small number of differentially expressed genes (Figure 3A and Table S2-S7), consistent with limited differences identified in our complete *CHD3* knockout versus wild-type differential gene expression analysis on bulk transcriptomic data from whole day-50 organoids (Figure S12 and Table S8). For the ventral cell types, we detected just 3-5 differentially expressed genes, likely due to the relatively small number of cells with a ventral cell type annotation in the original single cell transcriptomics dataset (as few as 164/4506 = 3.6% of cells in 010A^WT/WT^ C3; Figure 3A and Figure S11B). We therefore focused on the dorsal cell types (Figure 3A-B and Figure S13), with cortical excitatory neurons having the highest *CHD3* expression in wild-type organoids (Figure 2D, Figure S7). For cortical intermediate progenitors the expression of the differentially expressed genes only explained the difference between the wild-type and homozygous knockout condition (Figure S13B). However, for cortical neural progenitors and excitatory neurons, the differentially expressed genes also separately clustered two of the three 010A^WT/KO^ samples (Figure 3C, Figure S13B), and when plotting the normalized counts for a selection of these differentially expressed genes, we observed a dosage-response in expression level for most of them based on *CHD3* genotype (Figure 3D and Figure S13C).

**Figure 3.**
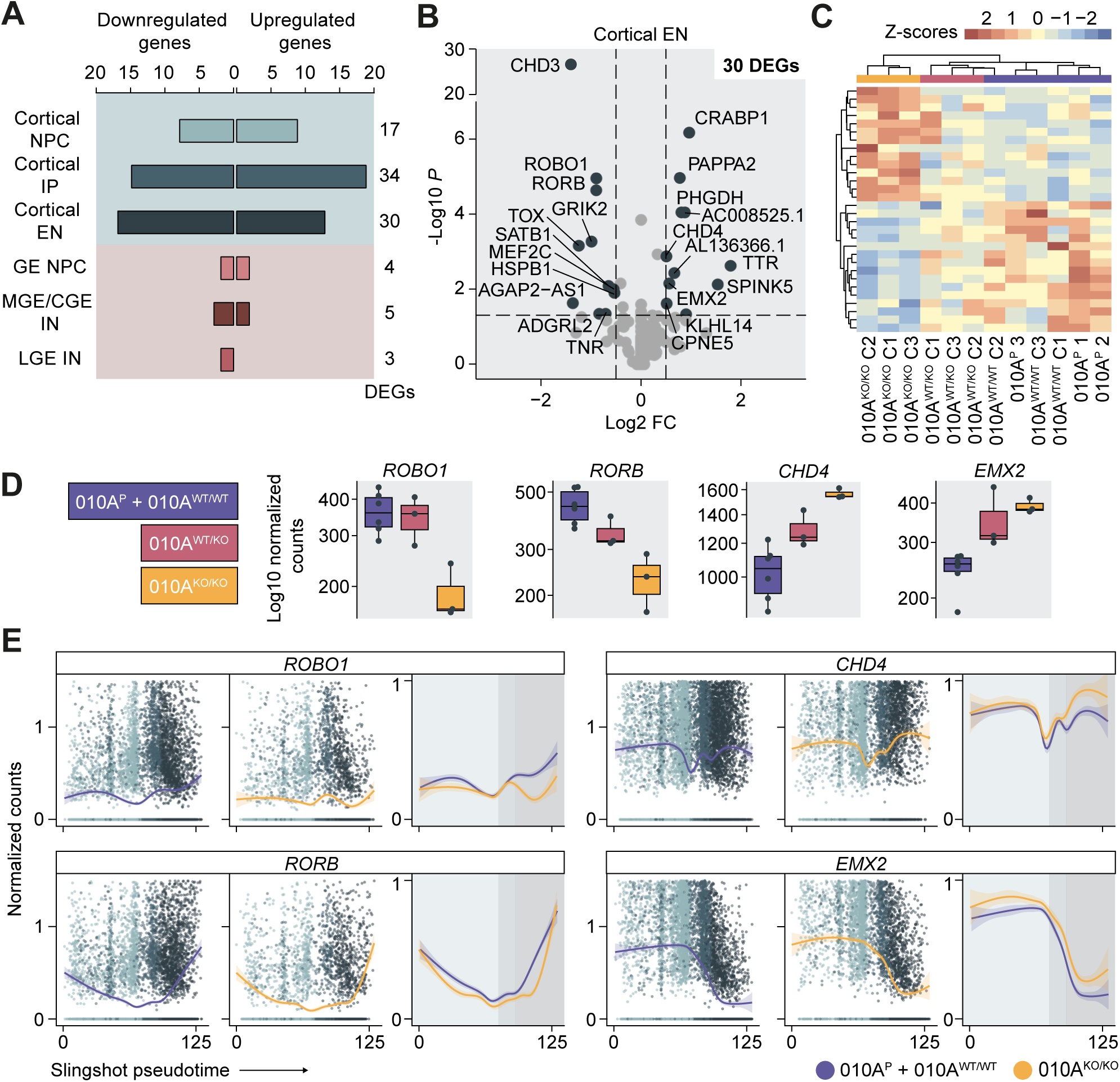
The effects of *CHD3* disruption on gene expression in cortical excitatory neurons in day-57 unguided neural organoids. **A**) Overview of the number of differentially expressed genes (DEGs with a *p* value < 0.05, 010A^KO/KO^ versus 010A^P^ + 010A^WT/WT^ comparison) in pseudobulk data for each dorsal and ventral cell type derived from the scRNA-sequencing dataset. **B**) Volcano plot of cortical excitatory neurons with the significant differentially expressed genes (*p* value < 0.05 and 0.5 < log2 fold change < 0.5) shaded in dark blue. **C**) A clustered heatmap based on the scaled expression values (Z-scores) of the significant differentially expressed genes in cortical excitatory neurons across each pseudobulk sample. **D**) Box plots of the pseudobulk normalized count data of excitatory neurons for a selection of differentially expressed genes across *CHD3* genotypes. The 010A^P^ + 010A^WT/WT^ condition is shown in purple, the 010AWT/KO condition in red, and the 010A^KO/KO^ condition in yellow. **E**) Scatter plots of the normalized counts from the scRNA-sequencing data for each cell over Slingshot pseudotimes for a selection of differentially expressed genes. The cortical neural progenitor cells are shown in light blue, the cortical intermediate progenitors in blue, and the cortical excitatory neurons in dark blue. The lines represent a smoothed fit ± standard error through the count values, with in purple the 010A^P^ + 010A^WT/WT^ and in yellow the 010A^KO/KO^ condition.

A number of genes differentially expressed in cortical excitatory neurons have well-established functions related to neurodevelopment. *ROBO1*, part of a receptor family important for axon guidance and cell adhesion^27^, and previously implicated in reading and language phenotypes^28^, was downregulated (Figure 3B and 3D-E). *RORB*, a marker of layer-V neurons and described to be important for the organization of pre- and post-synaptic organization in the barrel cortex of mice^29^ had a reduced expression as well. Moreover, *GRIK2 (GluR6)* and *MEF2C*, both downregulated in 010A^KO/KO^ cortical excitatory neurons (Figure 3B, Table S4), have roles related to synaptic function^30,31^ and have both been associated with neurodevelopmental disorders (MIM 619580 and 613443, respectively)^32,33^. Conversely, *EMX2 a*nd *CHD4*, normally highly expressed in radial glial cells^12,34^, were upregulated in response to loss of *CHD3* (Figure 3B and 3D-E). The increased *CHD4* expression could be a compensation mechanism for the *CHD3* disruptions, and may potentially partly underlie the subtle effects observed on organoid development after complete abrogation of *CHD3* expression. Overall, given that cell type composition in 010A^KO/KO^ organoids is unaffected, and that the limited number of downregulated genes seem related to more mature neuronal development and functions (also evident from the enriched gene ontology terms for the differentially expressed genes, including ‘neuron to neuron synapse’, ‘regulation of synaptic plasticity’ and ‘regulation of synapse organization’; Figure S14), CHD3 may not be essential for the switch from neural progenitor toward post-mitotic neuron, but rather plays roles in maturation and function of cortical neurons.

### CHD3 regulates other chromatin remodelers and acts upstream of SLIT and ROBO genes

To identify genes directly regulated by CHD3, we performed ChIP-sequencing on wild-type day-50 organoids. From the 20,426 non-redundant peaks across three replicates, 7,427 were called as high-confidence primary peaks (Figure 4A, Table S9). These peaks concentrated around transcription start sites (Figure 4B) and largely overlapped with promoter areas (Figure 4C). After gene annotation, we found enrichment of the Reactome pathway ontology terms of ‘chromatin modifying enzymes’ and ‘chromatin organization’, as well as gene ontology terms related to transcription factor binding and activity (Figure 4D-E and Figure S15A-B), suggesting that CHD3 controls the expression of other chromatin remodelers and transcriptional regulators. Strikingly and consistent with the findings from our differential gene expression analyses on the pseudobulk datasets, genes belonging to the ontology terms ‘signaling of ROBO receptors’ and ‘regulation of SLITs and ROBOs’ showed significant enrichment too (Figure 4D-E), as well as the KEGG pathway term ‘axon guidance’ (Figure S15C), providing further evidence of roles for CHD3 in axon and synapse development, guidance and growth. Moreover, the other significantly enriched ontology terms ‘translation’, ‘response of EIF2AK4 (GCN2)’ (Figure 4D), ‘Golgi-to-ER trafficking’ (Figure S15A) and ‘endocytosis’ (Figure S15C) are all processes that have been implicated in synaptic plasticity and neocortical development^35–38^, and ‘nonsense−mediated decay’ has been described to play a role in axon growth as well^39^. Indeed, the identified genes in these ontologies showed overlap with the ones from the ‘signaling of ROBO receptors’ and ‘regulation of SLITs and ROBOs’ ontology terms (Figure 4E), and converge on a role of CHD3 in axon development and function.

**Figure 4.**
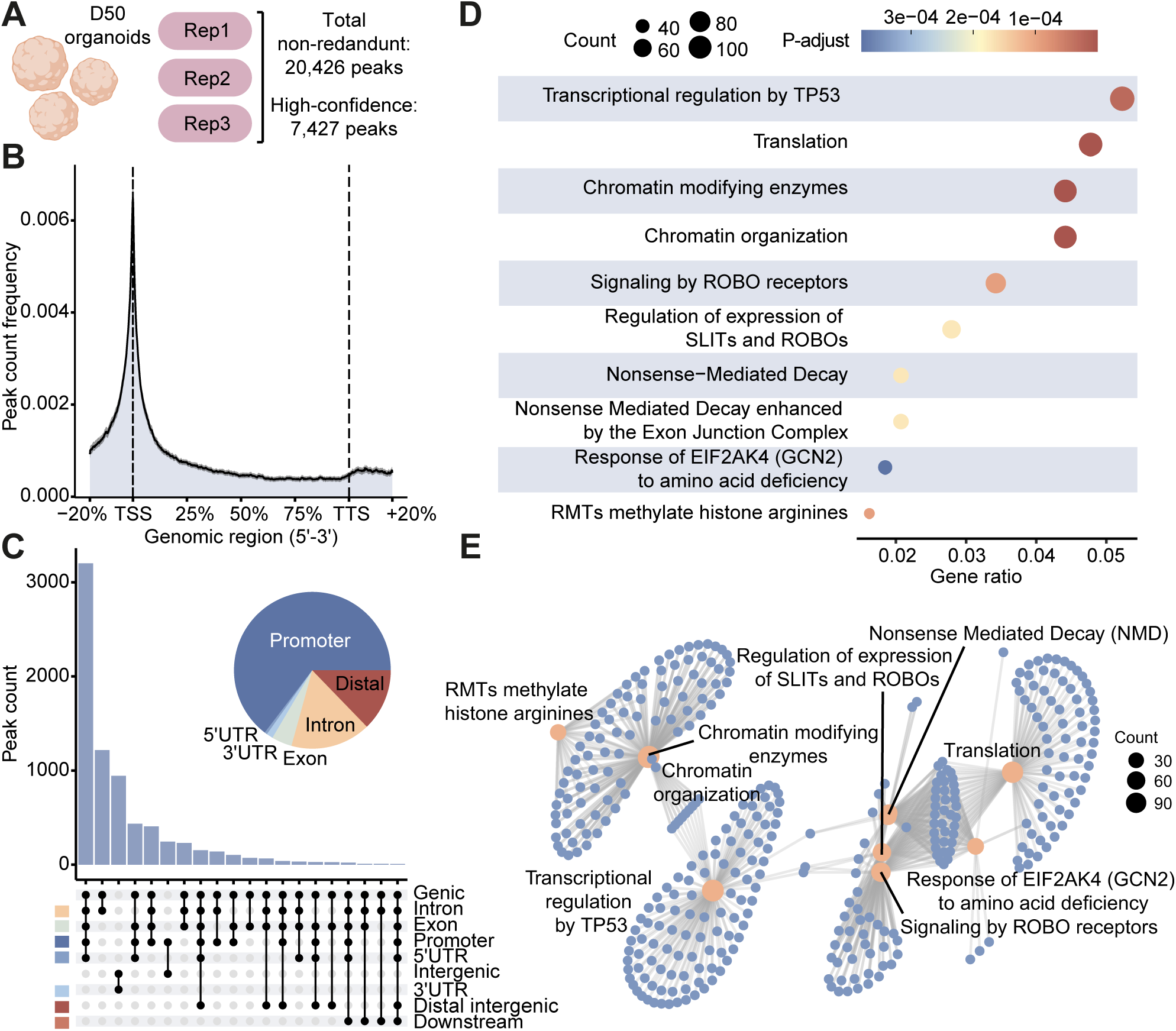
CHD3 primarily binds to promoter regions and the CHD3 binding sites are enriched for genes associated with chromatin organization and SLIT/ROBO functioning. **A**) Overview of the CHD3 ChIP-sequencing experiment, resulting in 7,427 high-confidence peaks. **B**) A binding-density profile of CHD3 across gene bodies ± 95% confidence interval, showing high coverage around the transcription start sites (TSS). **C**) An UpSet plot visualizing the intersections of CHD3 peaks across genomic annotations, with the bars showing the total peak count across these intersections. The pie chart visualizes the proportions of genomic annotations of CHD3 peaks, with promoter regions in dark blue, 5’ UTR regions in blue, 3’ UTR regions in light blue, exonic regions in green, intronic regions in yellow, and distal intergenic regions in red. **D**) A dotplot showing the results of a Reactome pathway enrichment analysis on the genes associated with CHD3 ChIP-peaks with a promoter annotation (4,799 out of 7,427 peaks). The colouring from blue to red represents the adjusted *p* value, and the dot size the number of genes mapping to the Reactome ontology term. **E**) A cnetplot depicting the connections between genes associated with the enriched Reactome ontology terms from (**D**).

### Genes differentially expressed upon *CHD3* knockout in stem cell-derived forebrain neurons imply roles in axon and synapse development

To validate our findings from neural organoids, particularly from the cortical excitatory neurons, we used a different independent culturing method to differentiate the 010A^P^, 010A^WT/WT^ C1-C3, 010A^WT/KO^ C1-C3 and 010A^KO/KO^ C2-C3 iPSC lines via NPCs (Figure S16) into day-21 forebrain neurons (Figure 5A). (Note that although the 010A^KO/KO^ C1 line generated good quality NPCs (Figure S16), the line did not differentiate well into forebrain neurons and consequently could not be employed for this part of the study.) In the two-dimensional forebrain neuronal cell model, CHD3 protein expression levels increased ∼25-30 fold in wild-type day-7 neural precursors and day-21 forebrain neurons when compared to wild-type iPSCs (Figure 5B and 5C), consistent with the relatively high levels of CHD3 expression observed in mature neuronal cell types in our neural organoids (Figure 2D and Figure S7 and S8). The decrease in CHD3-levels observed from day-7 neural precursors to day-21 forebrain neurons may be caused by differences in the amount of starting material, method of sample collection, as well as the semi-quantitative character of immunoblotting (Figure 5B and 5C)^40^. Independent of the *CHD3* genotype, all the differentiated cell lines gave rise to neurons positive for the somatodendritic marker MAP2 (Figure 5D), and we did not observe significant differences between conditions at protein level for pre-synaptic marker SYP (Synaptophysin 1) and post-synaptic marker PSD-95 (Figure S17).

**Figure 5.**
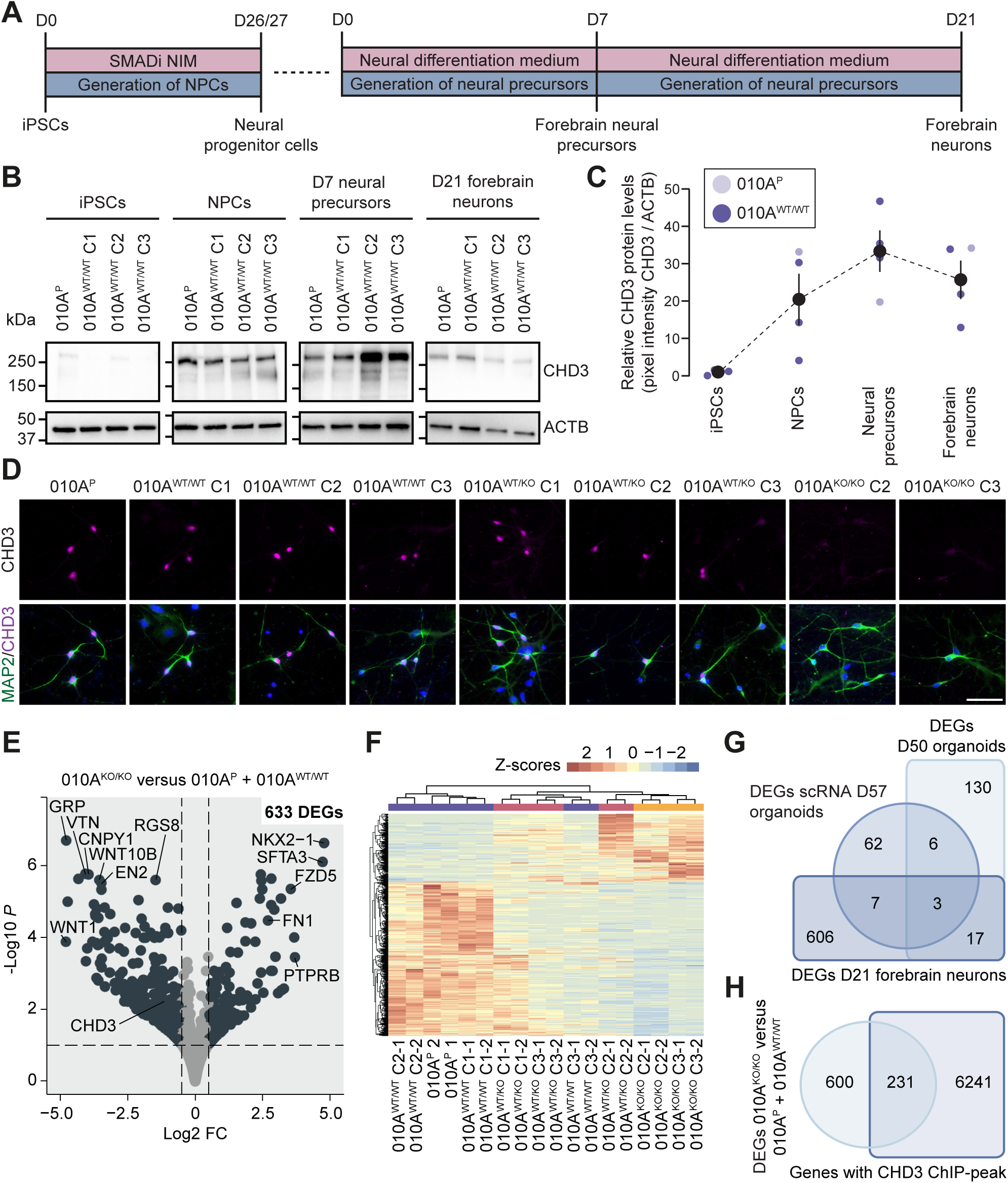
The effects of *CHD3* disruption on gene expression in day-21 stem cell-derived forebrain neurons. **A**) A schematic of the protocol used to generate NPCs and subsequently forebrain neurons, described in detail in the Methods section. **B**) Immunoblot of whole-cell lysates of iPSCs, NPCs, day-7 neural precursors and day-21 forebrain neurons of wild-type conditions for CHD3 protein. Expected molecular weight is ∼226 kDa. The blot was probed for ACTB to ensure equal protein loading. **C**) Quantification of the immunoblots shown in (**B**). **D**) Immunohistochemistry micrographs of day-21 forebrain neurons stained for neuronal marker MAP2 (green) and CHD3 (magenta). Nuclei were stained with Hoechst 33342. Scale bar = 50 μm. **E**) Volcano plot of day-21 forebrain neurons with the significant differentially expressed genes (*p* value < 0.05 and 0.5 < log2 fold change < 0.5) shaded in dark blue. **F**) A clustered heatmap based on the scaled expression values (Z-scores) of the significant differentially expressed genes in forebrain neurons across each sample. **G**) Venn-diagram showing the overlap between the differentially expressed genes (DEGs) identified in the transcriptomics datasets generated in this study. **H**) Venn-diagram showing the overlap between significant differentially expressed genes identified in this study upon complete *CHD3* knockout and genes associated with high-confidence CHD3 ChIP-peaks.

When we performed bulk RNA sequencing on the day-21 forebrain neurons, followed by differential gene expression analysis comparing 010A^KO/KO^ to wild-type neurons, we observed differences between these conditions, identifying 633 differentially expressed genes (Figure 5E and 5F, Table S10). Plotting the counts for a selection of differentially expressed genes, we again observed *CHD3* genotype-dependent dosage effects (Figure S18A). While only a small number of differentially expressed genes directly overlapped with the transcriptomic datasets independently generated from neural organoids (4.3%, 27/633; Figure 5G), we found ‘synaptic signaling’ to be one of the top enriched gene ontology terms, and identified ‘postsynaptic membrane’ and ‘axon guidance’ among the other enriched terms (Figure S18B), demonstrating functional convergence on synaptic development and function across the different neural organoid and forebrain neuron datasets (Figure 4D and Figure S14, S15 and S18).

Although we found evidence for an upregulation of CHD4 in forebrain neurons after CHD3 knockout in a small number of samples at protein level (Figure S17), in accordance with upregulation at transcript level in day-50 whole organoids and cortical excitatory neurons in day-57 organoids, *CHD4* was not among the significantly differentially expressed genes in the forebrain neurons according to bulk RNA screening.

When we overlapped all differentially expressed genes associated with a complete knockout of *CHD3* identified in this study with genes linked to CHD3 ChIP-peaks, we found about 28% (231/831 genes) to be present in these complementary datasets (Figure 5H, Table S11), suggesting that these 231 genes could be direct targets of CHD3-NuRD during early brain development. From those, *GAS8* and *ZHFX4-AS1* are examples that we found dysregulated upon disruption of *CHD3* in two of the three transcriptomic datasets (Figure S12 and S13, Table S11), and their genetic loci contained significant CHD3 ChIP-peaks located in an intronic enhancer and the promoter region respectively (Figure S19), making them especially strong candidates for regulation by this chromatin remodeler.

## Discussion

Using a combination of CRISPR-Cas9 gene-editing, investigations of independent cellular models of neurodevelopment, and subsequent transcriptomic and chromatin immunoprecipitation assays, we show that the expression of *CHD3* is particularly high in mature neurons, that this chromatin remodeling factor controls the expression of genes involved in axon development and guidance, and that its ablation affects the expression of a relatively small selection of genes associated with synapse formation, organization and function.

Unlike the profound effects of *Chd3* and *Chd5* knockdowns in murine models of brain development on the radial migration of cells out of the ventricular zone and the differentiation and specification of mature cortical neurons^12,17^, demonstrating a potential switch in the function of the NuRD-complex by replacing the position of Chd4^12^, we only detected subtle differences when we fully disrupted *CHD3* in human cellular models. In mice, Chd4-NuRD was found to promote the expression of *Sox2*, *Pax6* and *Tbr2/Eomes*, while Chd3-NuRD was reported to directly bind to these genes and repress their expression, thereby inducing the cell cycle exit of NPCs and initiating neuronal differentiation^12^. We indeed identified primary CHD3 ChIP-peaks in day-50 organoids in the genetic loci of *SOX2*, *PAX6* and *TBR2/EOMES*, but we did not find evidence for differences in their expression levels in the neural organoids, nor did we observe overt defects in radial migration in the developing cortical wall of the rosette-like structures or changes in the relative numbers of neural progenitors and mature neurons upon *CHD3* knockout.

Rather than a direct role for CHD3 in molecular layer specification of neurons as observed in mice, the functions of this protein in our human neuronal models seemed to instead converge on synaptogenesis, synapse function and axon guidance. Beyond the possibility of species-related differences in expression patterns and functions of mouse and human orthologues, which has been described before for various genes acting during neurodevelopment^41^, differences in the techniques and models used could potentially explain discrepancies between the earlier studies and the present one. In particular, while 2-month neural organoids are a good model to study the transition of NPCs from their proliferation to differentiation state, they do not fully capture the development of neurons into distinct layer identities^42^, and neither do the stem cell-derived forebrain neurons. Effects that are specific for subpopulations of cortical neurons are challenging to identify in such cellular models. However, although the thus far identified CHD3-associated developmental processes in animal and human model systems may appear distinct, the establishment of cell polarity, radial migration, formation and organization of synapses, and subsequent specification of neuronal cell identity are all highly interdependent processes^43^, and so our results are not necessarily in disagreement with prior studies.

A possibility for the lack of overt effects of *CHD3* disruption in neural organoids on neuronal differentiation capacities could be compensatory mechanisms via *CHD4*. Our transcriptomic analyses identified upregulation of *CHD4* expression in neural organoids fully lacking *CHD3* expression, specifically in cortical excitatory neurons that normally express *CHD3* at relatively the highest levels. Although prior work has underscored the non-redundant functions of CHD3 and CHD4 proteins^12,16^, both chromatin remodelers are actually highly conserved (71.62% amino acid identity between them) and closest to each other within the CHD family of proteins. Indeed, in human cell lines CHD3 and CHD4 share many interaction partners and regulate distinct but also shared target genes^16^. Analyzing the available data on CHD4-NuRD target genes^14,15^, we found overlaps (albeit modest) with our CHD3 ChIP-seq annotated genes (Table S12). Moreover, a subset of the differentially expressed genes in our CHD3 knockout neuronal models overlap with the Chd4-NuRD described targets in granule cells of the developing cerebellar cortex, including CPNE6/7^14^, COX6B2 and CRABP1^15^. This overlap shows that there may be a certain level of redundancy between CHD4 and CHD3 within a neurodevelopmental context and/or that CHD4 may be able to partly take over from CHD3. In previously reported work, the disruption of *Chd4* in mouse thymus^44^ and satellite cells from mouse skeletal muscles^45^ induced compensatory upregulation of *Chd3* as well. Overexpression of CHD3 has been reported to result in a decrease of *CHD4* transcript levels in human cell lines^16^. A potential compensation mechanism might maintain the availability of (alternative) NuRD complexes, rescuing crucial functions of the complex. It remains unclear whether the upregulation of *CHD4* involves a disrupted inhibitory feedback loop of class-II CHDs onto each other. Moreover, with the current analyses we do not know if a CHD4-NuRD complex can indeed mitigate the loss of CHD3 expression and subsequently the loss of CHD3-NuRD complexes. Based on our data, it would be interesting in future studies to examine CHD4 expression levels in cases of CHD3-associated disorder, in particular in individuals with loss-of-function variants.

The large majority of *de novo CHD3* variants identified in individuals with a neurodevelopmental disorder are missense variants^3,8,9^. Heterozygous *CHD3* variants with a predicted loss-of-function effect have mostly been identified in familial cases, with variable expressivity and/or reduced penetrance as an underlying mechanism, suggesting that a second hit or additional mutational load may be required for a *CHD3* loss-of-function variant to result in a neurodevelopmental phenotype^10^. Moreover, to our knowledge, no cases have so far been reported of people carrying homozygous or compound heterozygous loss-of-function variants in this gene. Hence, our genetically engineered full *CHD3* knockout cell lines are not modelling the genetic condition of individuals with CHD3-associated disorder, but rather help to uncover fundamental roles of CHD3 during early human neurodevelopment. By providing insights into potential functions of this gene and the pathways that it regulates, these neuronal cell models can enhance our understanding of disorder, especially since some of the downstream effects of a complete disruption may overlap with pathways disturbed in CHD3-associated disorder.

Taken together, using a gene knockout approach in human cellular models of cortical development, we show that, although *CHD3* expression levels strongly increase once NPCs start differentiating, CHD3 is not a crucial factor for inducing neuronal differentiation. Instead, CHD3 may play roles in supporting and defining the growth of axons, and formation and development of synapses, ultimately contributing to the generation of correctly specified and functional mature neurons. Consistent with our results, a recent study focusing on epigenetic factors that control neuronal maturation identified CHD3 as one of the chromatin remodelers crucial for controlling the pace of maturation of cortical neurons^46^. To better understand these potential functions of *CHD3*, in particular in guiding proper cell polarity and neuronal maturation and connectivity, future work should focus on the roles of the gene in models of neuronal networks and circuitry. Moreover, coupling transcriptomics and ChIP-seq to assays of DNA accessibility and histone modifications will further increase our understanding of how this important regulatory factor modifies molecular pathways to contribute to normal neuronal differentiation and maturation.

## Methods

### Cell line and cell culture

The BIONi010-A (K1P53) iPSC line (male, 15-19y, European Bank for induced pluripotent Stem Cells) derived from a healthy donor^23^, was cultured on plates coated with Matrigel (Corning) in mTeSR1 medium (Stem Cell Technologies) at 37 °C with 5% CO_2_. Medium was replaced daily and cells were passaged using Versene solution (Gibco) when confluency of 70-80% was reached. The chromosomal integrity of the cell line was confirmed by Cell Guidance Systems with an aCGH array before initiating CRISPR-Cas9 experiments.

### Gene editing with CRISPR-Cas9

In order to target exon 3 of the human *CHD3* gene, present in all three major isoforms (NM_005852.3, NM_001005273.2 and NM_001005271.2), we designed a guide RNA using CRISPOR^47^ with a high specificity (0.97), a high predicted efficiency (0.62-0.70) and a small number of predicted off-targets (19 off-targets, Table S1): 5’-AATATGGAACCGGACCGGGT *CGG*-3’. BIONi010-A cells were pre-treated with Y-27632 (10 μM; Selleckchem) for 30 min, disassociated using TrypLE Express (Gibco), and passaged through a 40 μm strainer to obtain a single-cell suspension. The guide was delivered as an Alt-R CRISPR-Cas9 sgRNA (IDT) after forming a protein complex with the Alt-R *Streptococcus pyogenes* Cas9 Nuclease V3 (IDT), using the P3 Primary Cell 4D-Nucleofector TM X kit (Lonza Biosciences) in combination with the Alt-R Cas9 Electroporation Enhancer (IDT). The electroporation was performed with an AMAXA 4D CoreUnit (CA137 program; Lonza Biosciences). Cells were maintained in mTeSR1 medium supplemented with 10 μM Y-27632 for 4-6 days and afterwards passaged one time to recover from the electroporation. To isolate colonies derived from single cells, cells were disassociated with TrypLE, passaged through a 40 μm strainer and seeded at a low density in Matrigel-coated 100 mm dishes in mTeSR1 supplemented with 1x CloneR (Stem Cell Technologies). After 7-8 days, iPSC colonies were manually picked and transferred to a 96-well plate in medium supplemented with 1x CloneR until ready to passage. Clones were split to three wells, to prepare cryovials for freezing in mFreSR (Stem Cell Technologies) and to isolate genomic DNA using Squishing buffer (10 mM Tris-HCl (pH 8.0), 1 mM EDTA, 25 mM NaCl, 1:5 1 mg/ml Proteinase K) to screen for the introduced mutations and CRISPR-Cas9 off-target effects.

### Screening of CRISPR-Cas9 edited cell lines

The iPSC clones were screened for mutations introduced by CRISPR-Cas9 gene editing in exon 3 of the *CHD3* gene by amplifying the target region of the guide RNA using PCR. For each clone, a PCR reaction was prepared with isolated gDNA, Phusion Green Hot Start II High-Fidelity PCR Master Mix (Thermo Fisher) and primers annealing to the target region (Table S13). PCR products were sent for Sanger sequencing (Eurofins Scientific) and the resulting Sanger traces were analyzed using the ICE CRISPR analysis tool^48^ to identify heterozygous and compound heterozygous or homozygous out-of-frame mutations. Positive clones were selected for expansion and further characterization. To assess off-target effects, five off-targets were selected for screening with PCR followed by Sanger sequencing (Table S1; primers used are described in Table S13). All selected CRISPR-Cas9 edited clones underwent molecular karyotyping using the KaryoStat HD Assay (Thermo Fisher).

### Organoid differentiation

Neural organoids were cultured using a well-established protocol as previously described^20,21^, with minor adjustments. Single-cell suspensions were prepared using TrypLE, and 9,000 cells were seeded per well in an ultra-low attachment U-bottom 96-well plate (Corning) in mTeSR1 medium supplemented with 50 μM Y-27632. The day of seeding was considered day 0 (Figure 2D). Half of the medium was replaced every other day. On day 5, neural induction was started by changing the medium to neural induction medium: DMEM/F12, 1x N2, 1x GlutaMAX (all Gibco), 1x minimum essential media-nonessential amino acids (MEM-NEAA) and 1:1000 1 mg/ml heparin from porcine intestinal mucosa (both Sigma). At day 12, embryoid bodies were transferred to drops of 20 μL of ice-cold Matrigel, and the Matrigel was allowed to solidify at 37 °C for > 30 min. Afterwards the embedded embryoid bodies were transferred to 60 mm dishes in differentiation medium: 50% DMEM/F12, 50% Neurobasal medium, 0.5x N2, 1x B27 minus VitA, 1x GlutaMAX, 0.5x MEM-NEAA, 50 μM 2-Mercaptoethanol, 1x Pen/Strep, 1:4000 human insulin (9.5-11.5 mg/mL; Sigma). The next day, the dishes were moved to a CO_2_ Resistant Shaker (Thermo Fisher), and organoids were cultured for the rest of the protocol under shaking conditions (40 rpm, orbit of 19 mm). Medium was replaced every other day. On day 18, the medium was changed to differentiation medium containing B27 with VitA (Gibco). On day 50, organoids were collected for RNA isolation and chromatin immunoprecipitation by pooling three organoids together and snap-freezing on dry-ice (for bulk RNA-seq and ChIP-seq), or fixed in 4% paraformaldehyde (Electron Microscopy Supplies Ltd) for 30 min at room temperature followed by 90 min at 4 °C for immunostainings. On day 57, four organoids were pooled and prepared for single cell RNA-seq. During the neural organoid differentiation, bright-field images were taken with an Axiovert A-1 microscope (Zeiss). To calculate the total area of organoids at early stages of the differentiation, a threshold was applied to the images followed by automated particle size analysis using ImageJ 1.51n software.

### Forebrain neuron differentiation

iPSCs were differentiated into neural progenitor cells using the STEMdiff SMADi Neural Induction Kit (Stem Cell Technologies). In brief, iPSCs were seeded at a density of 2.5 x 10^5^ cells/cm^2^ on Matrigel-coated plates in mTeSR1 medium supplemented with Y-27632. After 24 h the medium was changed to StemDiff Neural Induction medium supplemented with SMAD inhibitors, and the medium was replaced daily. Cells were differentiated in neural induction medium for 26-27 days and passaged using Accutase (Sigma) at high density onto Matrigel-coated plates for a total of three times during this period. After neural progenitor differentiation, neural progenitor cells were characterized and cryo-preserved. To generate forebrain neurons, the STEMdiff Forebrain Neuron Differentiation Kit (Stem Cell Technologies) was used. Briefly, cryo-vials of neural progenitor cells were thawed and cells were seeded at a density of 1.25 x 10^5^ cells/cm^2^ on Matrigel-coated dishes in StemDiff Neural Progenitor medium (Stem Cell Technologies) on day 0. The next day, day 1, medium was changed to StemDiff Forebrain Neuron Differentiation medium, and medium was replaced daily for seven days. At day 7, cells were passaged using Accutase, and seeded at a density of 2.0 x 10^4^ cells/cm^2^ on glass coverslips coated with Laminin/Poly-L-Ornithine (Sigma and Merck-Millipore), or at a density of 5.0 x 10^4^ cells/cm^2^ on Laminin-/Poly-L-Ornithine coated plates in StemDiff Forebrain Neuron Maturation medium (Stem Cell Technologies) supplemented with 1x Pen/Strep. Half medium changes were performed every other day. At day 21, the forebrain neurons on coverslips were fixed in 4% paraformaldehyde for 15 min at room temperature, and the cells cultured in plates were directly lysed in either 1x RIPA buffer (ThermoFisher) to obtain protein lysates or RLT buffer (Qiagen) for RNA isolation.

### Immunostainings of iPSCs, neural progenitors and forebrain neurons

iPSCs and neural progenitor cells were grown on Matrigel-coated coverslips and day-21 forebrain neurons on Laminin/PLO-coated coverslips. Cells were fixed with 4% paraformaldehyde for 15 min at room temperature. Then, cells were blocked and permeabilized for 1 h at room temperature with 5% horse or donkey serum (Vector or Sigma) and either 0.1% Triton-X100 for iPSCs and neural progenitors, and 0.025% Triton-X100 for forebrain neurons. Antibodies were diluted in blocking buffer (5% horse or donkey serum in PBS). Primary antibodies were incubated overnight at 4 °C and secondary antibodies for 1.5 h at room temperature. Nuclei were stained with Hoechst 33342 (Invitrogen) before cells were mounted in DAKO fluorescent mounting medium (DAKO). Primary antibodies: rabbit-anti-OCT4 (1:1000, AB19857, Abcam); mouse-anti-SSEA4 (1:500, AB16287, Abcam); goat-anti-SOX2 (1:500, af2018, R&D systems); mouse anti-TRA-1-60 (1:200, AB16288, Abcam); rabbit anti-PAX6 (1:500, GTX113241, GeneTex); mouse anti-SOX10 (1:100, sc-365692, Santa Cruz Biotechnology); mouse anti-MAP2 (1:2000, M2320, Sigma); rabbit anti-CHD3 (1:1000, ab109195, Abcam). Secondary antibodies: donkey anti-rabbit Alexa Fluor 488 (1:1000, Invitrogen, A21206); donkey anti-rabbit Alexa Fluor 594 (1:1000, Invitrogen, A21207); donkey anti-rabbit Alexa Fluor 647 (1:1000, Invitrogen, A31573); chicken anti-mouse Alexa Fluor 594 (1:1000, Invitrogen, A21201); donkey anti-goat Alexa Fluor 647 (1:250, Jackson Immuno Research, 705-605-147); donkey anti-mouse Alexa Fluor 488 (1:1000, Invitrogen, A21202). Fluorescence images were acquired with an Axiovert A-1 epifluorescence microscope or a LSM880 confocal microscope and ZEN Black Image Software (Zeiss).

### Immunostainings of neural organoids

Fixed day-50 and day-57 neural organoids were cryoprotected in 30% sucrose overnight at 4 °C, embedded in Neg-50 Frozen Section Medium (Thermo Fisher) and cryosectioned at 8 μm. Sections selected for immunostainings were rehydrated in PBS at room temperature for 20 min. Afterwards, sections were blocked and permeabilized with 5% horse or donkey serum and 0.25% Triton-X100 in PBS for 1 h at room temperature. Antibodies were diluted in blocking buffer (5% horse or donkey serum and 0.25% Triton-X100 in PBS). Primary antibodies were incubated overnight at 4 °C and secondary antibodies for 2 h at room temperature. Nuclei were stained with Hoechst 33342 (Invitrogen) before sections were mounted in DAKO fluorescent mounting medium (DAKO). Primary antibodies: mouse anti-PAX6 (1:500, 862002, BioLegend); sheep anti-EOMES/TBR2 (1:250, AF6166, R&D Systems); rabbit anti-CHD3 (1:500, ab109195, Abcam); chicken anti-TBR1 (1:500, AB2261, Millipore); rabbit anti-TBR1 (1:500, ab31940, Abcam); rat anti-BCL11B/CTIP2 (1:500; ab18465, Abcam); guinea pig anti-TUBB3 (1:250, 302 304, Synaptic Systems). Secondary antibodies: donkey anti-mouse Alexa Fluor 488 (1:1000, Invitrogen, A21202); donkey anti-mouse Alexa Fluor 594 (1:1000, Invitrogen, A21201); donkey anti-rabbit Alexa Fluor 594 (1:1000, Invitrogen, A21207); donkey anti-rabbit Alexa Fluor 647 (1:1000, Invitrogen, A31573); donkey anti-chicken Alexa Fluor 647 (1:250, Jackson Immuno Research, 703-605-155). Donkey anti-rat Alexa Fluor 488 (1:1000, Invitrogen, A21208); donkey anti-guinea pig Alexa Fluor 488 (1:250, Jackson Immuno Research, 706-545-148); donkey anti-sheep Alexa Fluor 647 (1:1000, Invitrogen, A21448). Fluorescence images were acquired with an AxioScan Z1 microscope and ZEN Blue Image Software (Zeiss).

### Immunoblotting

Whole-cell lysates were prepared by treating three pooled snap-frozen day-30 neural organoids, TrypLE-treated and pelleted iPSCs, Accutase-treated and pelleted 7-day neural precursor cells or day-21 forebrain neurons directly lysed in cell culture plates with 1x RIPA buffer (Thermo Fisher) supplemented with 1x PIC (cOmplete protease inhibitor cocktail, Roche) and 1% PMSF (Sigma). Cells were lysed for 20 min at 4 °C followed by centrifugation for 20 min at 12,000 rpm. Protein concentrations were measured in the cleared lysates using the Pierce BCA Protein Assay Kit (Thermo Fisher). Then, lysates were combined with Laemmli loading buffer (Bio-Rad) supplemented with 10% TCEP, boiled for 5 min at 95 °C and then loaded on 4–15% Mini-PROTEAN TGX Precast Gels (Bio-Rad). Gels were transferred onto polyvinylidene fluoride membranes using the Trans-Blot Turbo Transfer System (Bio-Rad), and membranes were blocked in 5% milk for 1 h at room temperature. After blocking, membranes were probed overnight at 4 °C with: rabbit anti-CHD3 antibody (1:1000; Abcam, ab109195); mouse anti-CHD4 (1:1000, Millipore, MABE455); chicken anti-SYP (1:1000, Synaptic Systems, 101006); mouse anti-PSD95 (1:1000, NeuroMab, 75-348); mouse anti-ACTB (Sigma, A5441). Next, membranes were incubated for 1.5 h at room temperature with 1:5000-1:10,000 HRP-conjugated antibodies: goat anti-rabbit (Jackson Immuno Research); goat-anti-mouse (Jackson Immuno Research); goat anti-chicken (Jackon Immuno Research, 103-035-155). The signal of ACTB was visualized with the Novex ECL Chemiluminescent Substrate Reagent Kit (Invitrogen) and all other targets with the SuperSignal West Femto Maximum Sensitivity Substrate Reagent Kit (Thermo Fisher) using a ChemiDoc XRS + System (Bio-Rad). Quantification of the signal intensity was done using ImageJ 1.51n software, measuring the inverted mean gray values of the detected bands, corrected for background signal and normalized to ACTB. Statistical analysis of the immunoblot quantification results was performed with GraphPad Prism 9 software using a one-way ANOVA followed by a Dunnett’s *post hoc* test. All original uncropped immunoblot images presented in this study are included in Figure S20.

### Single-cell RNA sequencing and analyses

Four day-56/day-57 organoids were pooled and cut in pieces using a sterile blade. The cut organoids were placed in pre-warmed Accutase supplemented with 1:2000 DNAseI and 1:2000 RNAase Inhibitor (both NEB), and slowly pipetted up and down using wide-bore pipette tips. After 30-60 minutes, cells were centrifuged for 5 min at 300x g, washed with PBS, and then passed through 30 and 20 μm filters (Miltenyi Biotec). The filtered cell suspension was centrifuged for 5 min at 300x g, and cells were resuspended in 500 μL differentiation medium. The resuspended cells were pipetted on top of a three-layered Percoll (Sigma) gradient which was centrifuged for 5 min at 300x g, to separate the cells from debris. The fraction that contained the cells was centrifuged 5 min at 300x g, and the cell pellet was resuspended in ice-cold PBS with 0.04% BSA. Cells were counted and cell viability was assessed based on Trypan Blue (BioRad) staining. Afterwards, ∼5000-7000 cells per sample were loaded on a Chromium Next GEM Chip G using the Chromium Next GEM Single Cell 3’ Kit v3.1 (both 10x Genomics). The dual-index library was prepared using the Library Construction Kit (10x Genomics), and quality control was performed with the RNA 6000 Nano Kit on a 2100 Bioanalyzer Instrument (both Agilent). Libraries were sent to Novogene for 150 bp paired-end sequencing on the Illumina Novaseq 6000 platform (400 million reads per sample, 120 Gb clean data per sample). Cellranger v3.1^49^ was used for demultiplexing of the data. Then, the count function from Cellranger v6.0.1^49^ was used to map the reads to the refdata-gex-GRCh38-2020-A. The generated raw feature matrices were loaded as individual SeuratObjects using the Seurat v4.3 package^50^, and subsequently combined into a single object using the Seurat merge function for further filtering. Genes with no counts, or that were only expressed in less than 10 cells were removed from the data. Additionally, only cells with > 500 UMIs, > 250 genes/features, a log10GenesPerUMI > 0.8, and a mitoRatio < 0.2 were kept. The data were then split again by sample, and each individual object was separately normalized. With the variance stabilizing transformation method, 2000 variable features were determined in each object and all samples were integrated into a single dataset based on the first 30 principal components. A reference dataset containing 46,977 cells from 2-month neural organoids^25^ (org2m_human_singlecells_GRCh38.rds, DOI:10.17632/z4jyxnx3vp.2, https://data.mendeley.com/datasets/z4jyxnx3vp/2) was used to identify transfer anchors and to subsequently annotate the integrated query dataset with the reference metadata^51^. The SPRING dimensionality reduction of the reference was used to visualize the cells of the annotated queried dataset. To perform a pseudotime analysis, cells with a dorsal cell type were extracted from the full dataset, and a principal component analysis and UMAP was run on this subset. The Slingshot v2.2.1 package^26^ was applied to infer a dorsal pseudotime trajectory, with the Cortical NPCs defined as starting cluster and the Cortical EN as end cluster.

### Pseudobulk differential gene expression analysis

To perform a pseudobulk differential gene expression analysis, the raw counts of the single-cell RNA sequencing dataset were extracted, and then aggregated across all individual cells for each sample per cell type. The aggregated count matrices were then loaded into DESeq2 1.34.0^52^ and normalized to perform differential gene expression analysis with ∼ genotype + batch + cell number as the design. Fold changes were shrunken using the ‘apeglm’ method^53^, and differentially expressed genes were filtered for an adjusted *p*-value <0.05 (FDR/Benjamini-Hochberg method).

### Bulk RNA-sequencing and analyses

RNA from day-50 neural organoids and day-21 forebrain neurons was sent to BGI (Hong Kong) for library preparation. Subsequently, RNA from the neural organoids underwent 150 bp paired-end stranded RNA sequencing on the BGI DNBseq platform. Due to lower input RNA, unstranded 100 bp paired-end sequencing was performed on the forebrain neuron samples. For the neural organoids, RNA isolation, library preparation and sequencing of the neural organoids was performed in two separate batches. First, the quality of the generated fastq files was assessed using MultiQC v1.11^54^. Then, the quasi-mapper Salmon v1.3.0^55^ was used to map the RNA-seq reads to the human reference transcriptome (Gencode release 45, GRCh38.p14) and to directly determine transcript abundance. The transcript abundance values were imported in DESeq2 v1.30.1^52^, which was then used to normalize these values across samples and to perform differential gene expression analysis. Principal component analysis was performed on regularized log transformed counts, and for the neural organoid data, batch normalization was done using the limma package^56^. The differential gene expression analysis was run with the ∼ genotype + batch design for neural organoids and the ∼ genotype design for forebrain neurons. Fold changes were shrunken using the ‘ashr’ method^57^, and differentially expressed genes were filtered for an adjusted *p*-value <0.5 (FDR/Benjamini-Hochberg method).

### ChIP-sequencing

Three samples of snap-frozen pooled wild-type day-50 neural organoids derived from the 010A^P^ line were sent to Active Motif (United States) for their FactorPath Service. The organoid tissue was fixed, and chromatin immunoprecipitation was performed on 40 μg chromatin with 10 μg rabbit anti-CHD3 antibody (Bethyl, A301-220A) according to Active Motif’s standard protocol. Illumina sequencing libraries were prepared from the chromatin immunoprecipitations and sequenced via single-end 75 bp sequencing. A pooled chromatin sample from the three independent samples was used as input control. Reads were aligned to the reference genome (hg38) using the BWA package^58^ by Active Motif. Next, the aligned reads were sorted and filtered with Samtools v1.11 to only keep aligned and unique reads. Duplicates, mitochondrial reads and reads that were not mapped to a known chromosome were removed. Reads mapping to genomic regions included in the ENCODE blacklist^59^ were removed as well. To ensure all samples had equal numbers of reads, all datasets were downsampled based on the sample with the lowest number of reads (∼34.5 million reads, downsampling ranged from 86.6 to 99.1% of total number of reads). Then, MACS v3.0.0b1 was used to call narrow peaks, with the pooled input sample as control. The ChIP-R v1.2.0 package^60^ was used to find primary peaks across all three replicate samples. These high-confidence peaks were loaded into ChIP-seeker v1.26.2^61^ to analyze peak coverage across genomic regions, and to perform peak annotation. The annotations were then used to perform functional enrichment analysis with the reactomePA v1.34^62^ and clusterProfile v3.18.1^63^ packages.

## Code availability

The computational code used in this study will be made available on GitHub upon publication.

Gene exon-intron schematics were generated in R 4.3.3 using ggplot2_3.5.0. The code has been made available as an R Shiny app for a range of different species (ExInPlotter): https://jho2024.shinyapps.io/ExInPlotter/

The following software was used for data analyses:

For bulk RNA-sequencing analysis we used Salmon v1.3.0 and R version 4.0.5 with the attached packages DESeq2_1.30.1, limma_3.46.0, pheatmap_1.0.12, tximport_1.18.0 and tximportData_1.18.0. For single cell RNA-sequencing analysis we used: Cellranger 6.0.1 and R version 4.1.2 with attached packages apeglm_1.16.0, DESeq2_1.34.0, pheatmap_1.0.12, Seurat_4.3.0, SingleCellExperiment_1.16.0 and slingshot_2.2.1 as well as R version 4.0.5 with attached package Seurat_4.0.2. For ChIP-sequencing analysis we used: ChIP-R 1.2.0, ChIPseeker_1.26.2, MACS 3.0.0b1 and Samtools 1.11, and R version 4.0.5 with attached packages clusterProfiler_3.18.1 and ReactomePA_1.34.0.

## Data availability

Single-cell RNA, bulk RNA and ChIP-sequencing data generated and analyzed in this study will be deposited for public access in the MPI for Psycholinguistics Archive (https://archive.mpi.nl/) upon publication.

## Acknowledgements

We would like to thank Dr. Karthikeyan Devaraju for sharing ideas and input at early stages of this study, Jula Peters for providing help and training setting up the neural organoid model, Arianna Vino for technical assistance, and Prof. Dr. Lisenka E.L.M Vissers for critical reading of earlier versions of the manuscript. This work was supported by the Max Planck Society.

