## Supplementary Figures for "The chromatin remodeler CHD3 is highly expressed in mature neurons and regulates genes involved in synaptic development and function"

### Affiliations

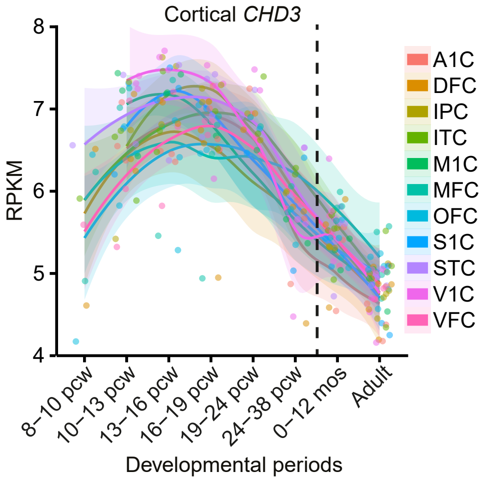


**Figure S1. *CHD3* expression in the human developing cortex.** Expression patterns of *CHD3* in different cortical regions based on the developmental human RNA-sequencing dataset of BrainSpan (http://www.brainspan.org/). The lines show loess curves fitted through the data points. A1C, primary auditory cortex; DFC, dorsolateral prefrontal cortex; IPC, posterior inferior parietal cortex; ITC, inferior temporal cortex; M1C, primary motor cortex; MFC, medial prefrontal cortex; OFC, orbital prefrontal cortex; S1C, primary somatosensory cortex; STC, superior temporal cortex; V1C, primary visual cortex; VFC, ventrolateral prefrontal cortex; pcw, post conception week.

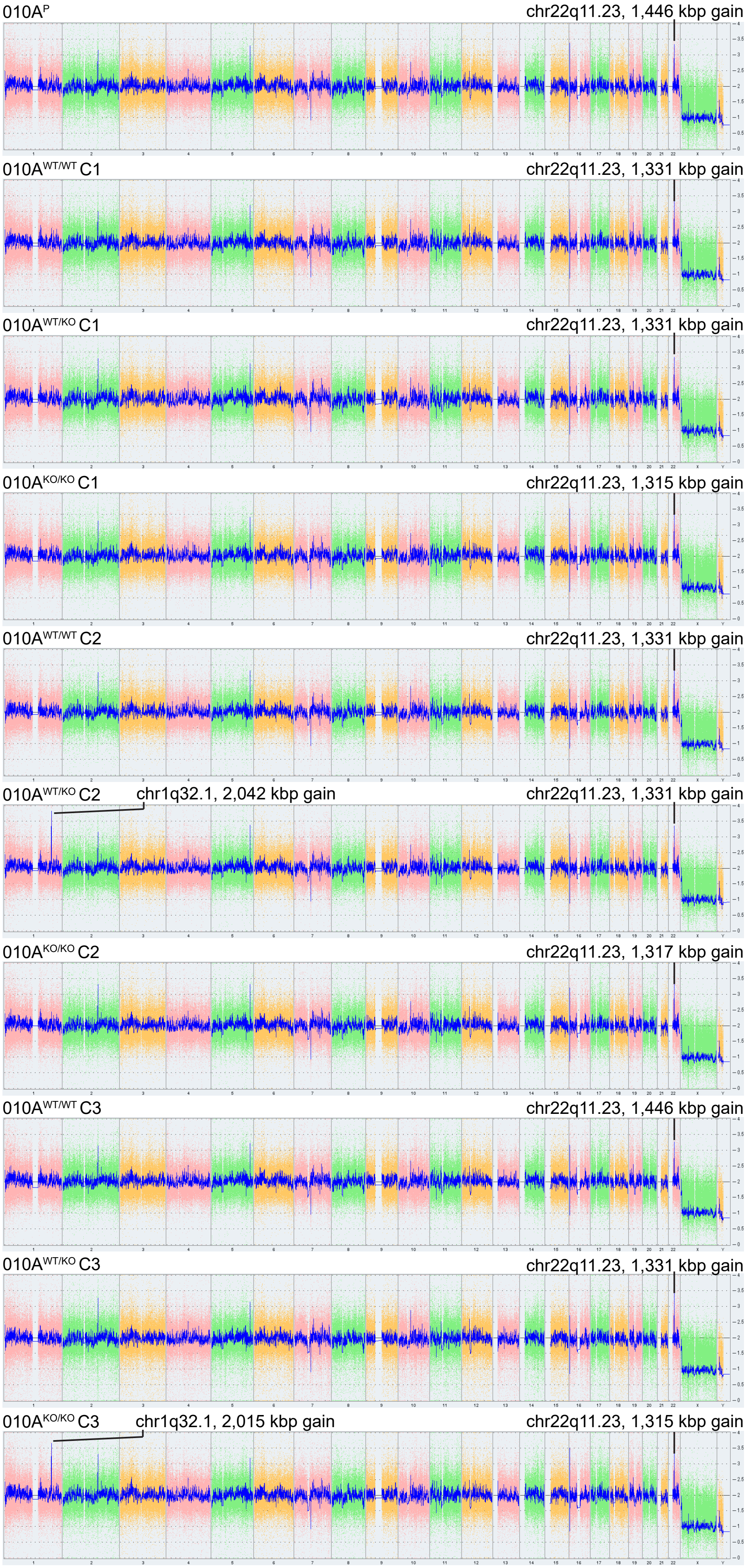


**Figure S2. Molecular karyotyping of CRISPR-Cas9 gene edited iPSC lines.** Molecular karyotyping was performed with the Cytoscan HT-CMA 96 array using the KaryoStat+ method. The size of structural aberrations that could be detected was >1 Mb for both chromosomal gains and chromosomal losses, although the resolution was depended on the location of the aberration in the chromosome. Due to a lower probe density on the telomere ends and centromeres, the resolution in those locations may have been closer to >5 Mb.

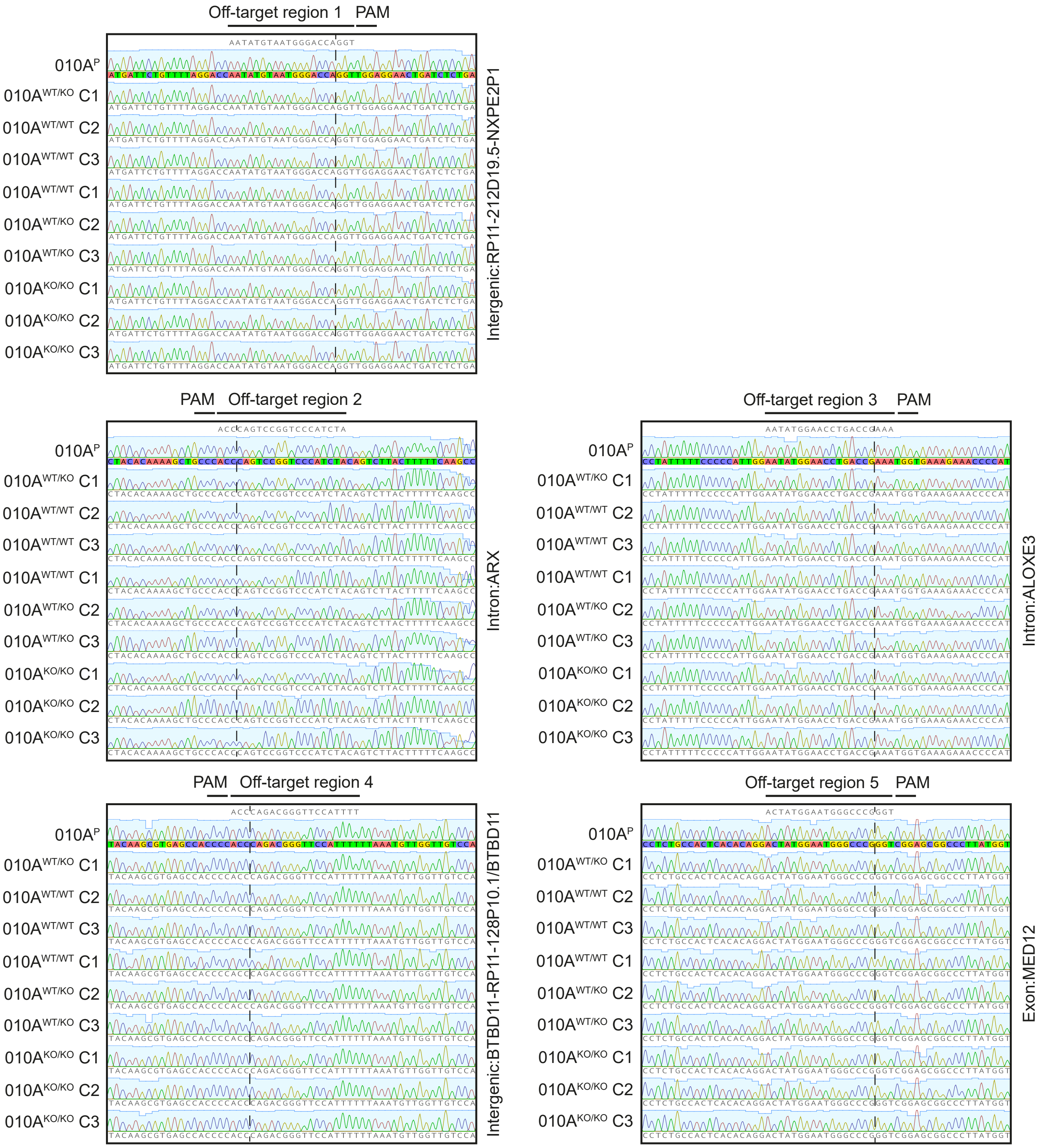


**Figure S3. CRISPR-Cas9 off-target analysis.** Five predicted off-targets for the CHD3 CRISPR-Cas9 gene editing design were selected (Table S1) for Sanger sequencing and subsequent analysis. Graphs show the Sanger traces for the off-target regions for each cell line, and the dashed lines indicate the predicted cut sites.

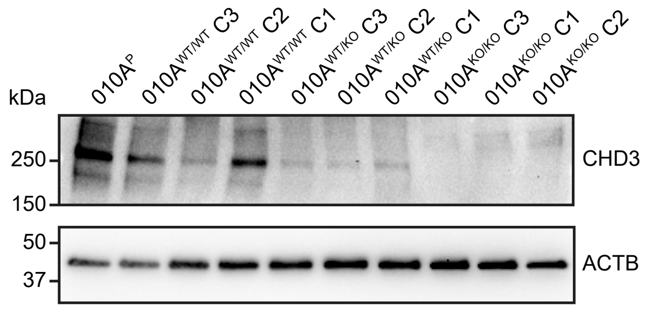


**Figure S4. Protein expression of CHD3 in iPSCs and the effect of the CRISPR-induced frameshift variants on CHD3 expression levels.** Immunoblot of whole-cell lysates of BiONi010-A iPSCs ($010A^{P}$) and the derived gene-edited cell lines ($010A^{WT/WT}$, $010A^{WT/KO}$ and $010A^{KO/KO}$) probed for CHD3. The expected molecular weight of CHD3 is ~226 kDa. The blot was also probed for ACTB to ensure equal protein loading.

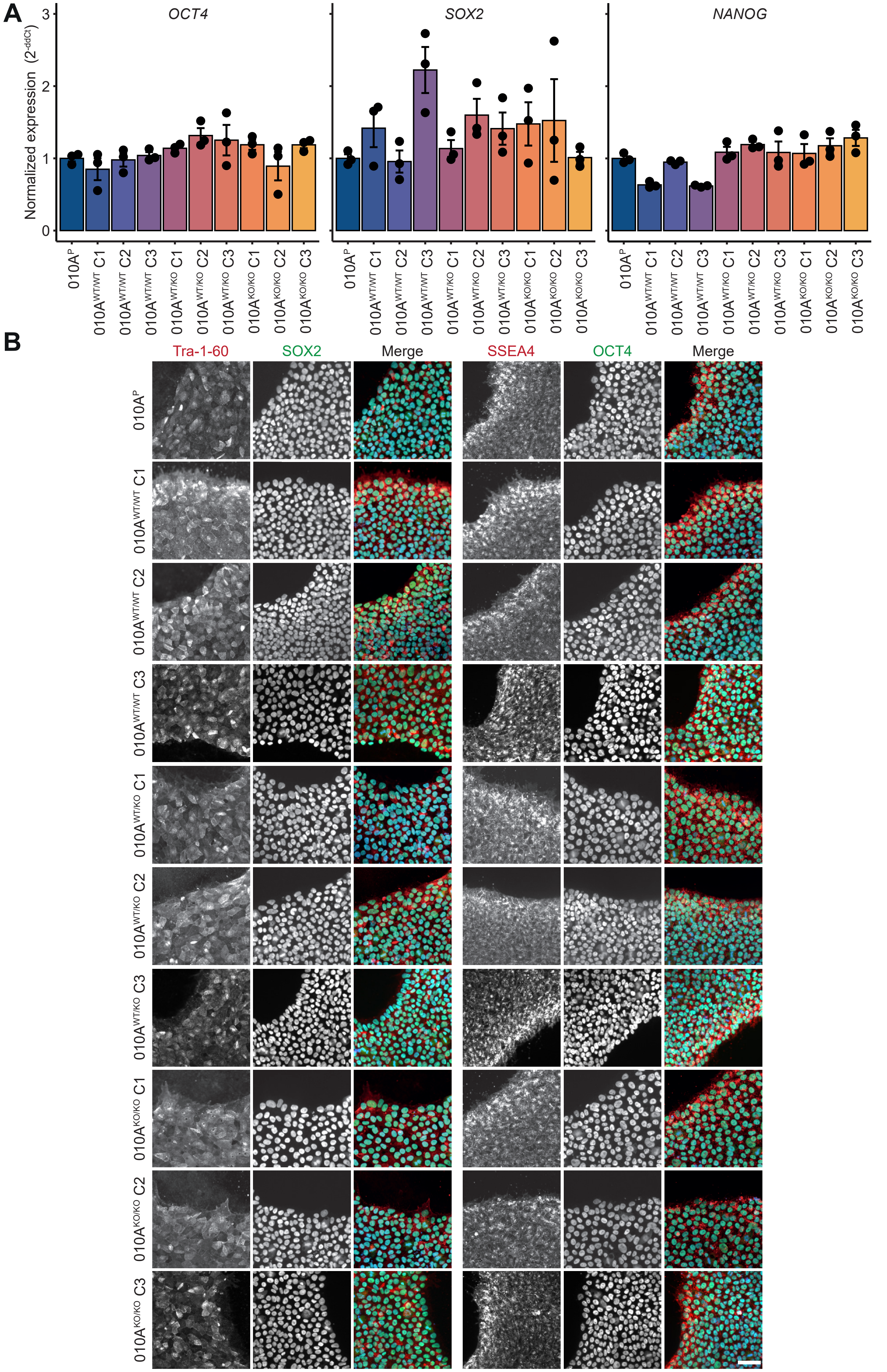


**Figure S5. Characterization of CRISPR-Cas9 gene-edited iPSC lines. A**) Bar plot representing the mean ± S.E.M transcript levels of pluripotency markers *OCT4*, *SOX2* and *NANOG* in iPSCs, assayed using qPCR (*n* = 3). **B**) Immunohistochemistry micrographs of iPSC colonies for pluripotency markers TRA1-60 (left, red), SOX2 (left, green), SSEA4 (right, red) and OCT4 (right, green). Nuclei were stained with Hoechst 33342 (blue). Scale bar = 50 µm.

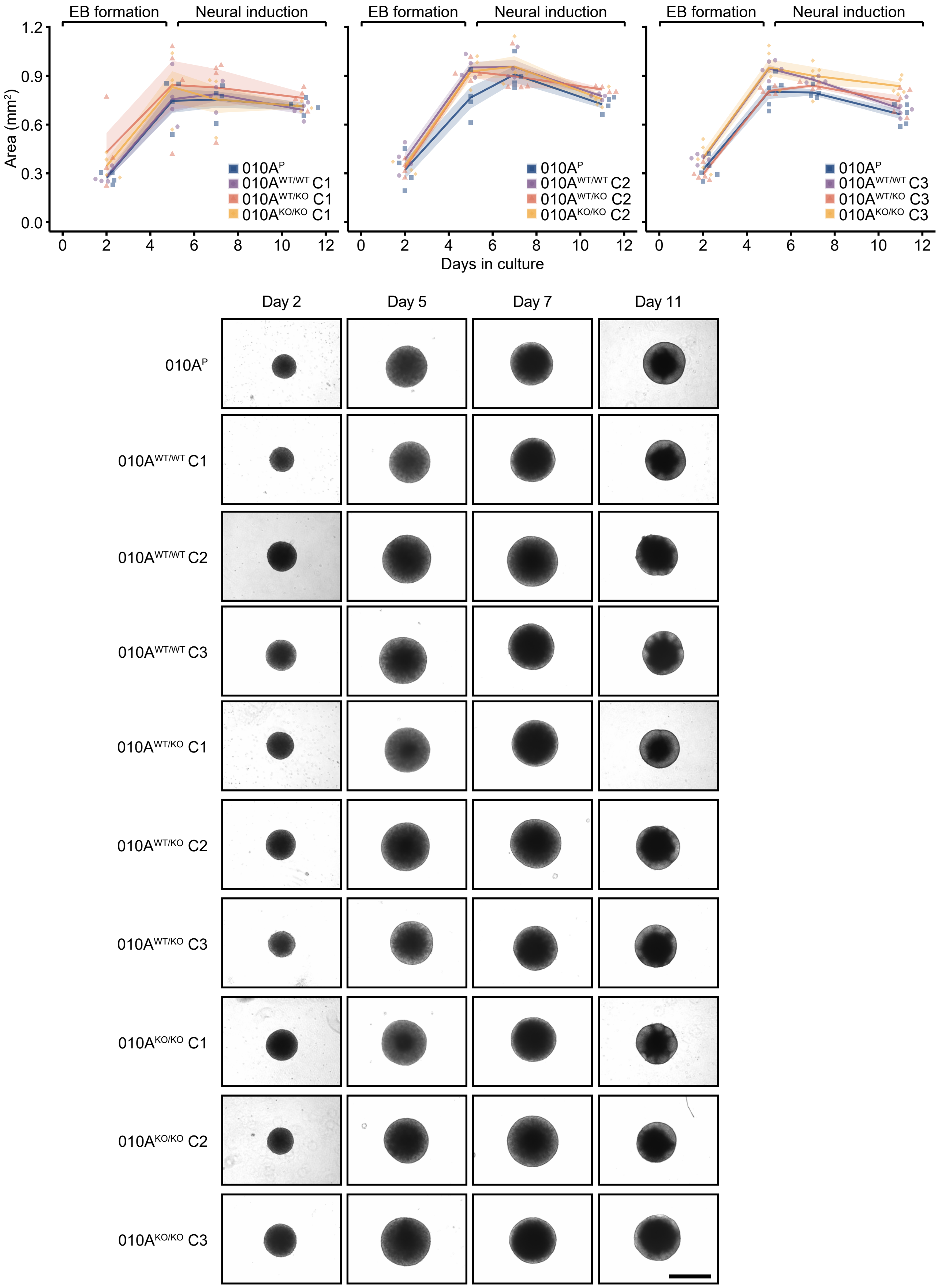


**Figure S6. Growth of embryoid bodies derived from CRISPR-Cas9 gene edited iPSC lines.** Top, a graph showing the surface area of embryoid bodies of each cell line at day 2, 5, 7 and 11 of the undirected neural organoid protocol. The lines connect the mean surface area at each day ± S.E.M. Bottom, representative bright-field images of day 2, 5, 7 and 11 unguided neural organoids for each cell line. Scale bar = 1000 µm.

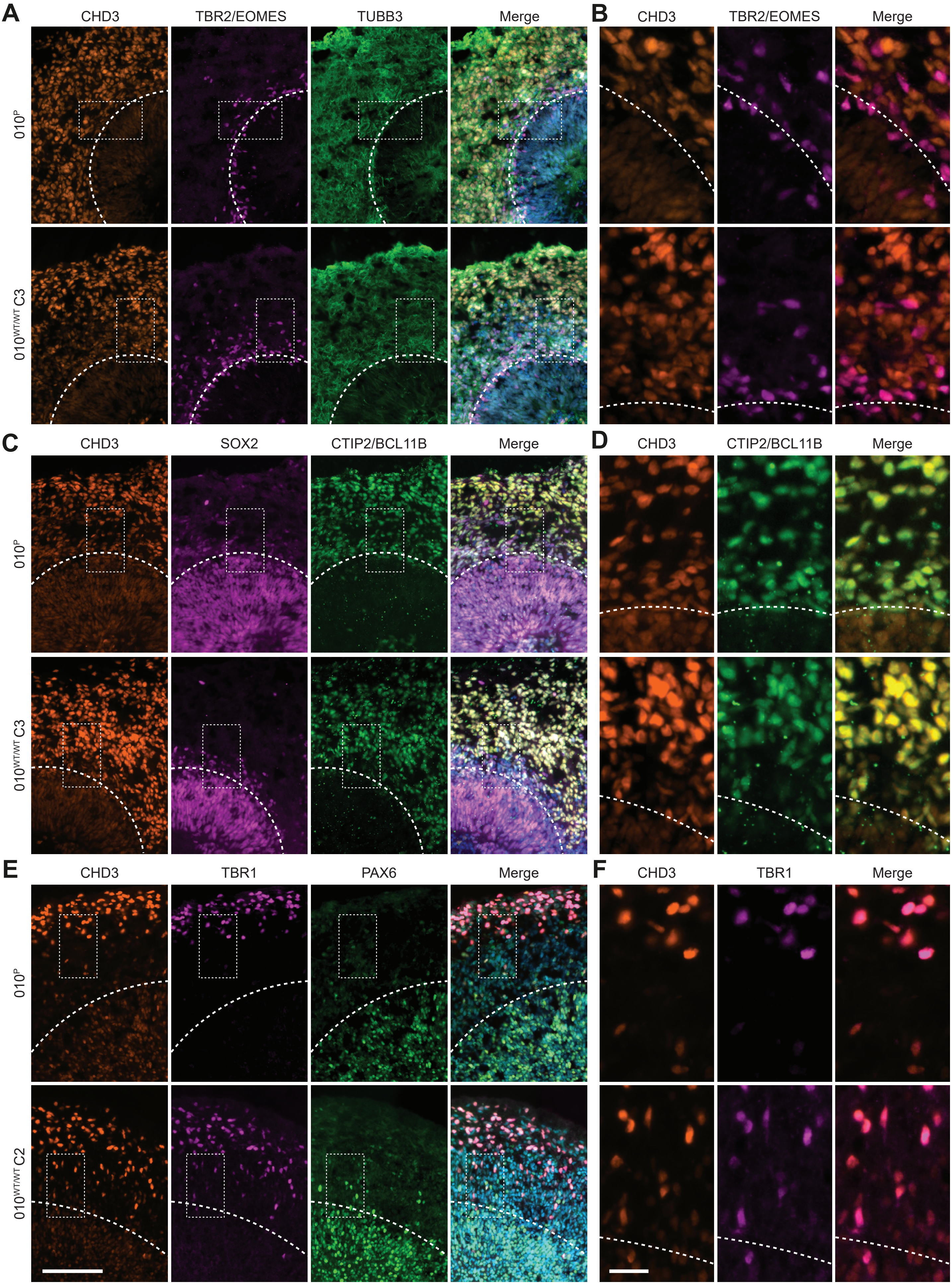


**Figure S7. Expression and localization of CHD3 in wild-type unguided neural organoids. A**) Micrographs of day-57 neural organoid rosettes showing CHD3-positive cells (orange), intermediate progenitor cells (TBR2/EOMES, magenta) and neuronal cells (TUBB3, green). The square shows a region of interest depicted in (**B**). **B**) Region of interest coloured as in (**A**). **C**) Micrographs stained for CHD3-positive cells (orange), neural progenitor cells (SOX2, magenta) and post-mitotic neurons (CTIP2/BCL11B, green). The square shows a region of interest depicted in (**D**). **D**) Region of interest coloured as in (**C**). **E**) Micrographs stained for CHD3-positive cells (orange), post-mitotic neurons (TBR1, magenta) and radial glia cells (PAX6, green). The square shows a region of interest depicted in (**F**). **E**) Region of interest coloured as in (**E**). **A-F**) The dashed white line marks the border of the ventricular zone. **A**, **C**, **E**) Nuclei are stained with Hoechst 33342. Scale bar = 100 µm. **B**, **D**, **F**) Scale bar = 20 µm.

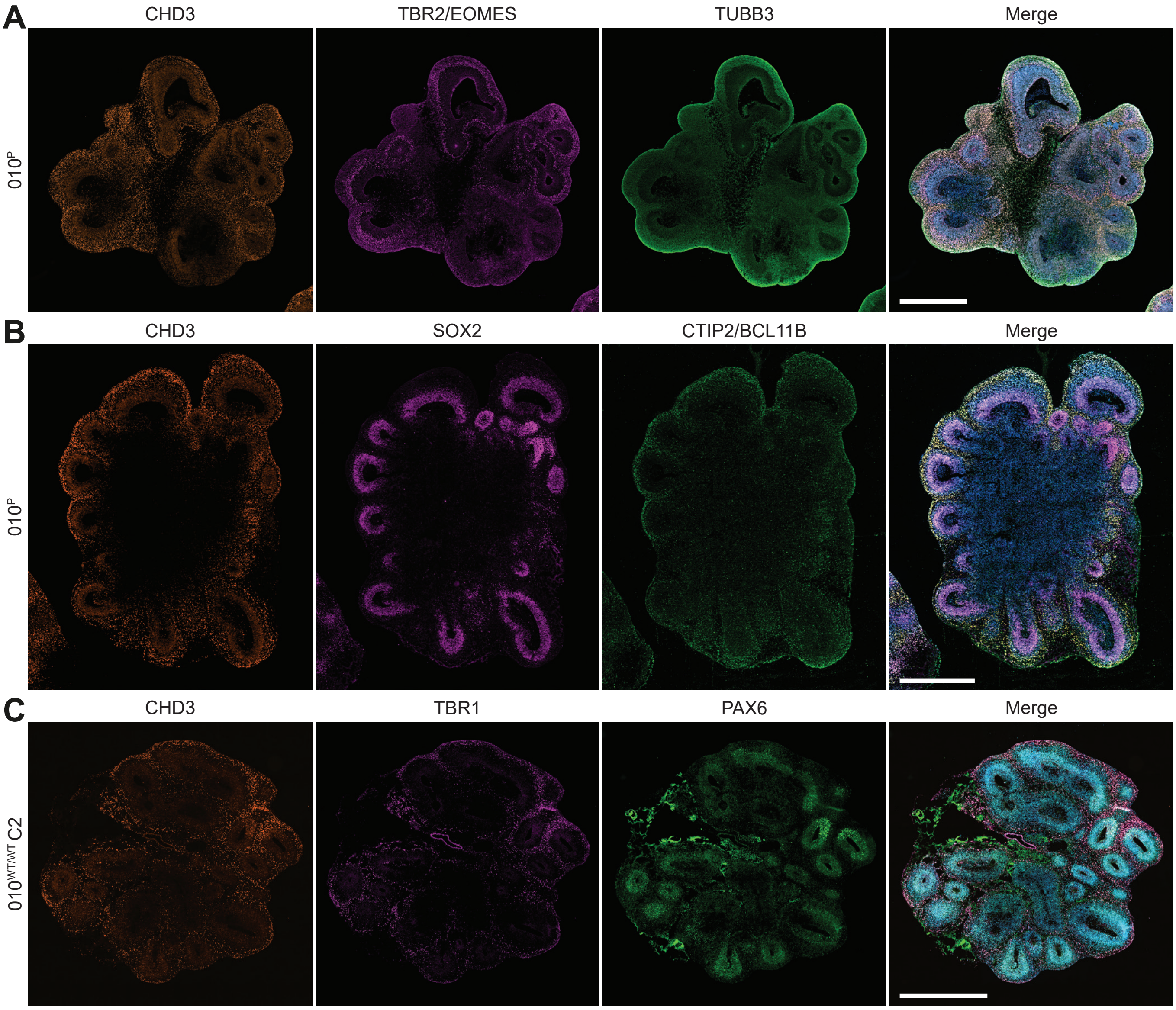


**Figure S8. Expression and localization of CHD3 in wild-type day-50 and day-57 unguided neural organoids. A**) Micrographs stained for CHD3-positive cells (orange), intermediate progenitor cells (TBR2/EOMES, magenta) and neuronal cells (TUBB3, green). **B**) Micrographs stained for CHD3-positive cells (orange), neural progenitor cells (SOX2, magenta) and post-mitotic neurons (CTIP2/BCL11B, green). **C**) Micrographs stained for CHD3-positive cells (orange), post-mitotic neurons (TBR1, magenta) and radial glia cells (PAX6, green). **A-C**) Nuclei are stained with Hoechst 33342. Scale bar = 1000 µm.

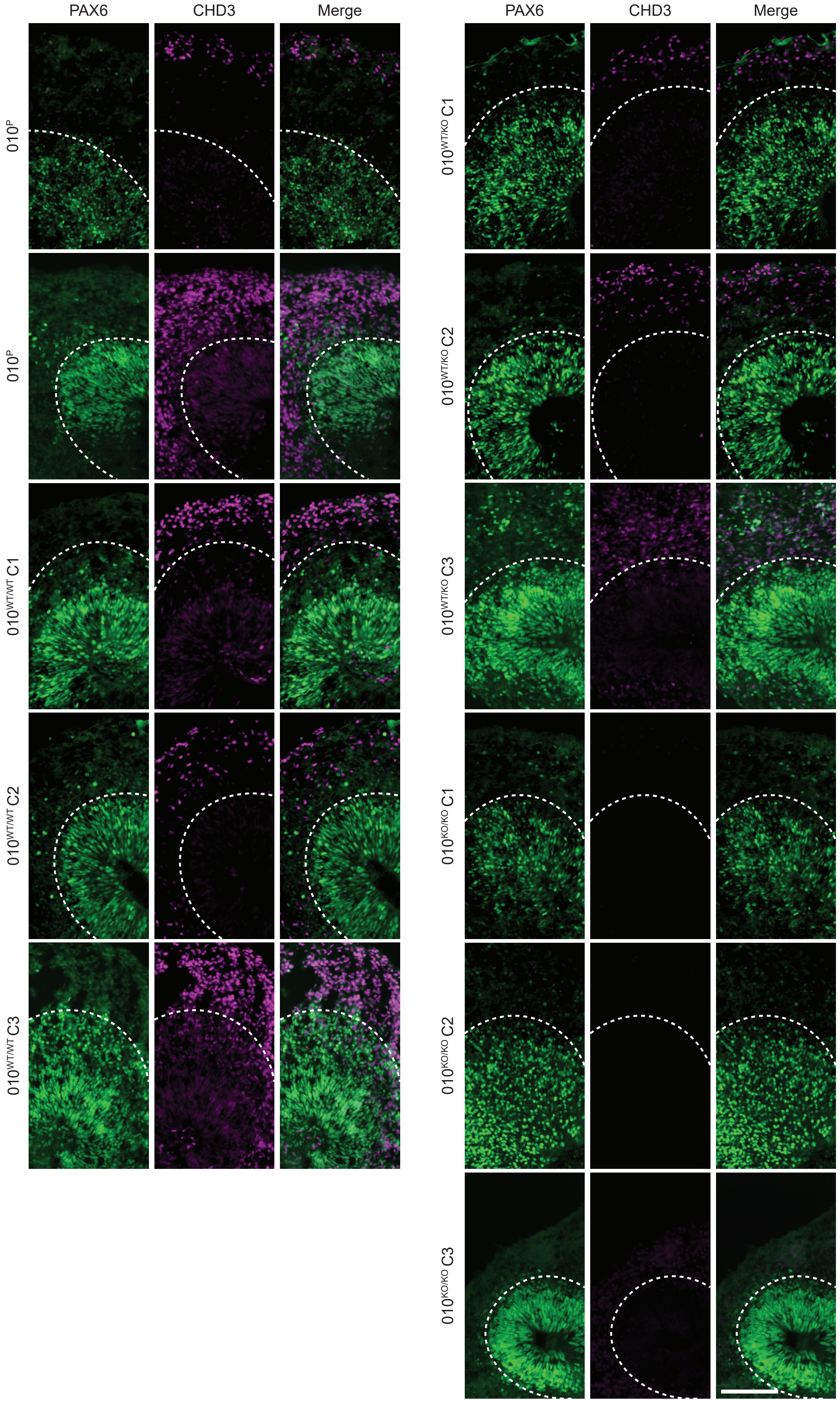


**Figure S9. CHD3 expression in unguided neural organoids derived from gene-edited iPSC lines.** Representative immunohistochemistry micrographs of day-57 organoid rosettes stained for radial glia cells (PAX6, green) and CHD3-positive cells (CHD3, magenta). Neural organoids grown from CHD3 full-knockout cell lines did not show any signal for CHD3. The dashed white line marks the border of the ventricular zone. Scale bar = 100 µm.

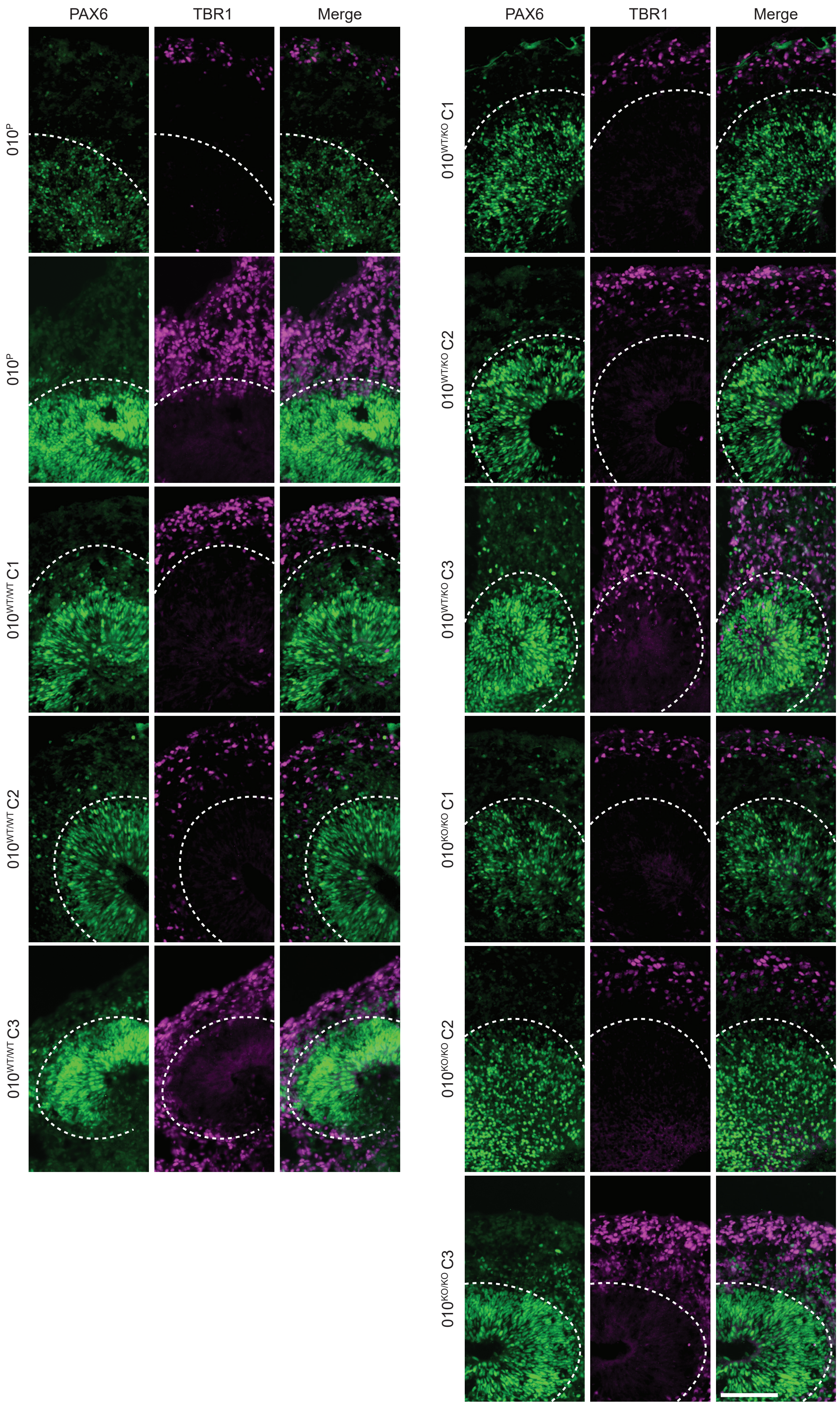


**Figure S10. Growth and structure of unguided neural organoids derived from gene-edited iPSC lines.** Representative immunohistochemistry micrographs of day-57 organoid rosettes stained for radial glia cells (PAX6, green) and post-mitotic neurons (TBR1, magenta). The dashed white line marks the border of the ventricular zone. The selected regions of interest are identical to the ones shown in Figure S9, except for the second region of $010^{P}$ and the selected regions of $010^{WT/WT}$ C3, $010^{WT/KO}$ C3 and $010^{KO/KO}$ C3. Neural organoids derived from all cell lines contain both PAX6- and TBR1-positive cells. Scale bar = 100 µm.

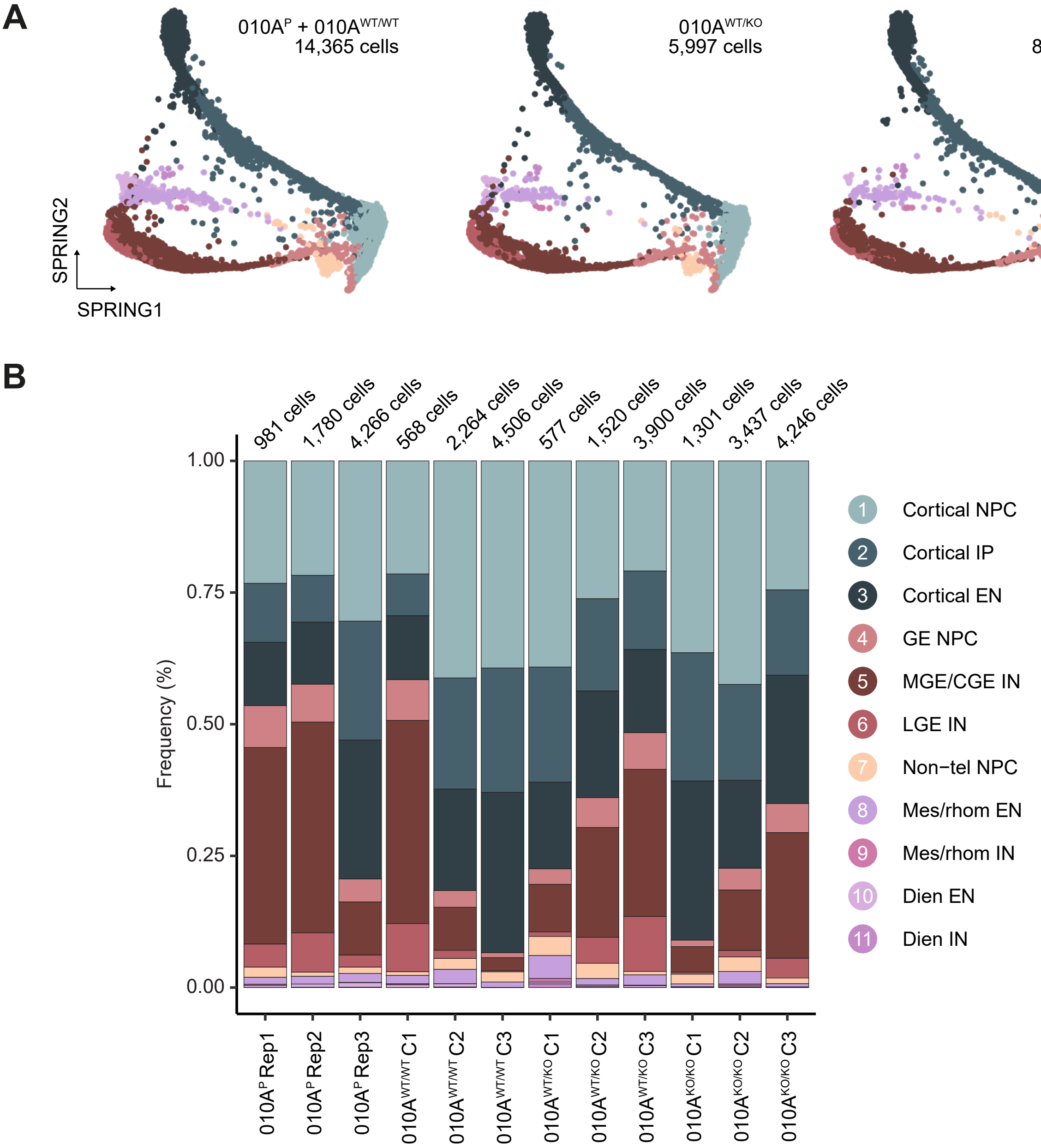


**Figure S11. The effects of *CHD3* disruption on cell composition of unguided neural organoids. A**) Scatter plots of the 29,346 cells from day-56/day-57 unguided neural organoids derived from the gene-edited 010A cell lines across the three *CHD3* genotypes. Cell types were annotated by querying the data on the dataset of Kanton et al. 2019, and cells were visualized in the same reduced dimensional space (SPRING). Colours as in Figure 2A. **B**) Stacked bar plots showing the relative cellular distribution for each sample. The number of cells per sample is indicated. CGE, caudal ganglionic eminence; EN, excitatory neuron; GE, ganglionic eminence; IN, inhibitory neuron; IP, intermediate progenitors; LGE, lateral ganglionic eminence; MGE, medial ganglionic eminence; NPC, neural progenitor cell.

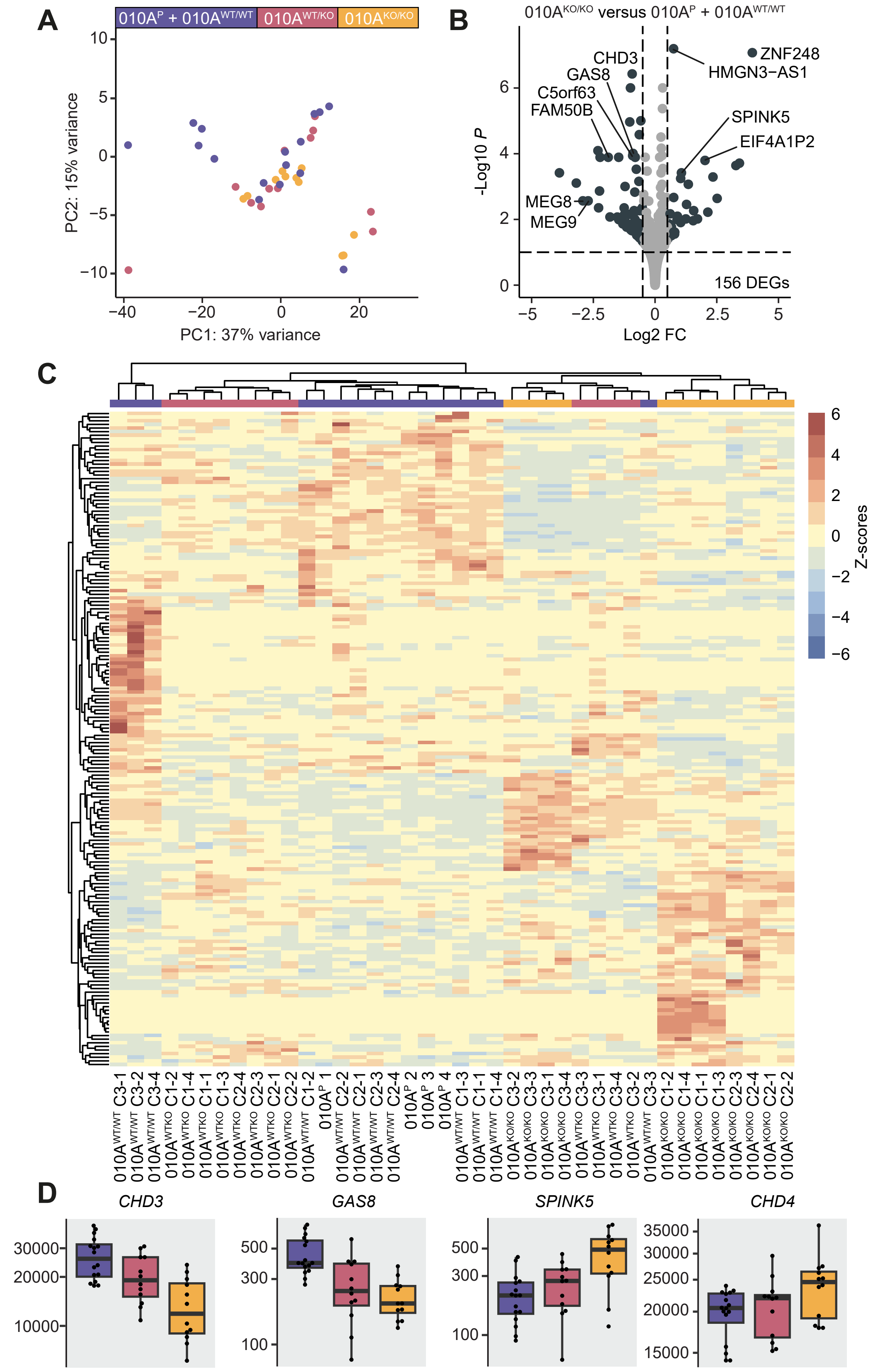


**Figure S12. The effects of *CHD3* disruption on gene expression in whole day-50 unguided neural organoids based on bulk RNA sequencing data. A**) Graph plotting principal component 1 (explaining 37% of the variance between the samples) and principal component 2 (explaining 15% of the variance between the samples), based on normalized counts. **B**) Vulcano plot with the significant differentially expressed genes (18 DEGs with *p* value < 0.05 and 0.5 < log2 fold change < 0.5) shaded in dark blue. In total 156 DEGs with *p* value < 0.05 were detected. **C**) A clustered heatmap based on the scaled expression values (Z-scores) of the significant differentially expressed genes across all samples. The $010A^{P}$ and $010A^{WT/WT}$ samples were shaded in purple, the $010A^{WT/KO}$ samples in red, and the $010A^{KO/KO}$ samples in yellow. **D**) Box plots of the bulk transcriptomics normalized count data of whole day-50 neural organoids for a selection of differentially expressed genes (*CHD3*, *GAS8*, *SPINK5* and *CHD4*) across *CHD3* genotypes. Colouring as in **C**.

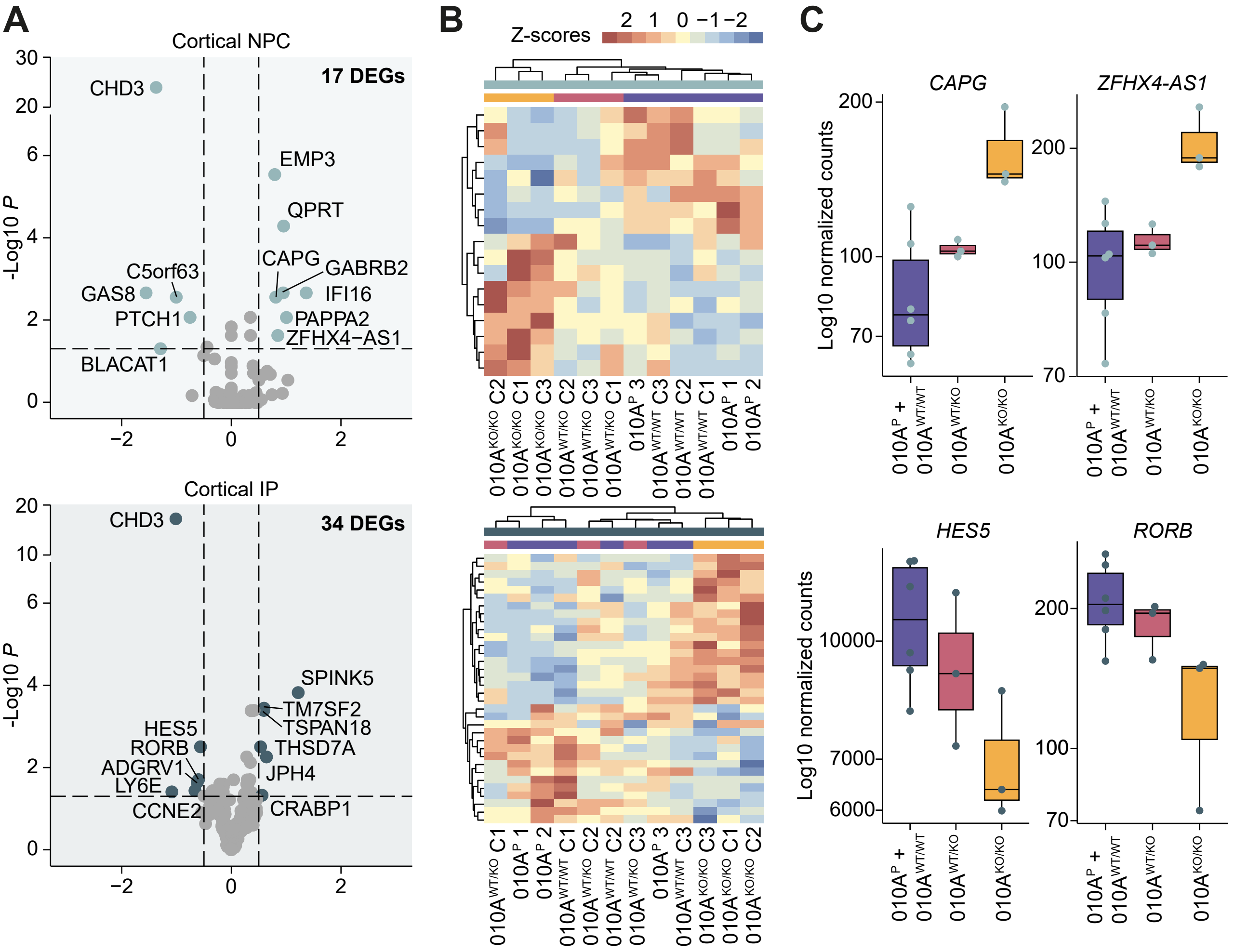


**Figure S13. The effects of *CHD3* disruption on gene expression in cortical neural progenitor cells and cortical intermediate progenitors in day-57 unguided neural organoids. A**) Vulcano plots of cortical neural progenitor cells (NPC; top) and cortical intermediate progenitors (IP; bottom) with the significant differentially expressed genes (*p* value < 0.05 and 0.5 < log2 fold change < 0.6) shaded in light blue and blue respectively. **B**) Clustered heatmap based on the scaled expression values (Z-scores) of the significant differentially expressed genes in cortical NPC (top) and cortical IP (bottom) across each pseudo-bulk sample. **C**) Box plots of the psuedo-bulk normalized count data of cortical NPC (top) and cortical IP (bottom) for a selection of differentially expressed genes across *CHD3* genotypes. The 010A$^{P}$ + 010A$^{WT/WT}$ condition is shown in purple, the 010A$^{WT/KO}$ condition in red and the 010A$^{KO/KO}$ condition in yellow.

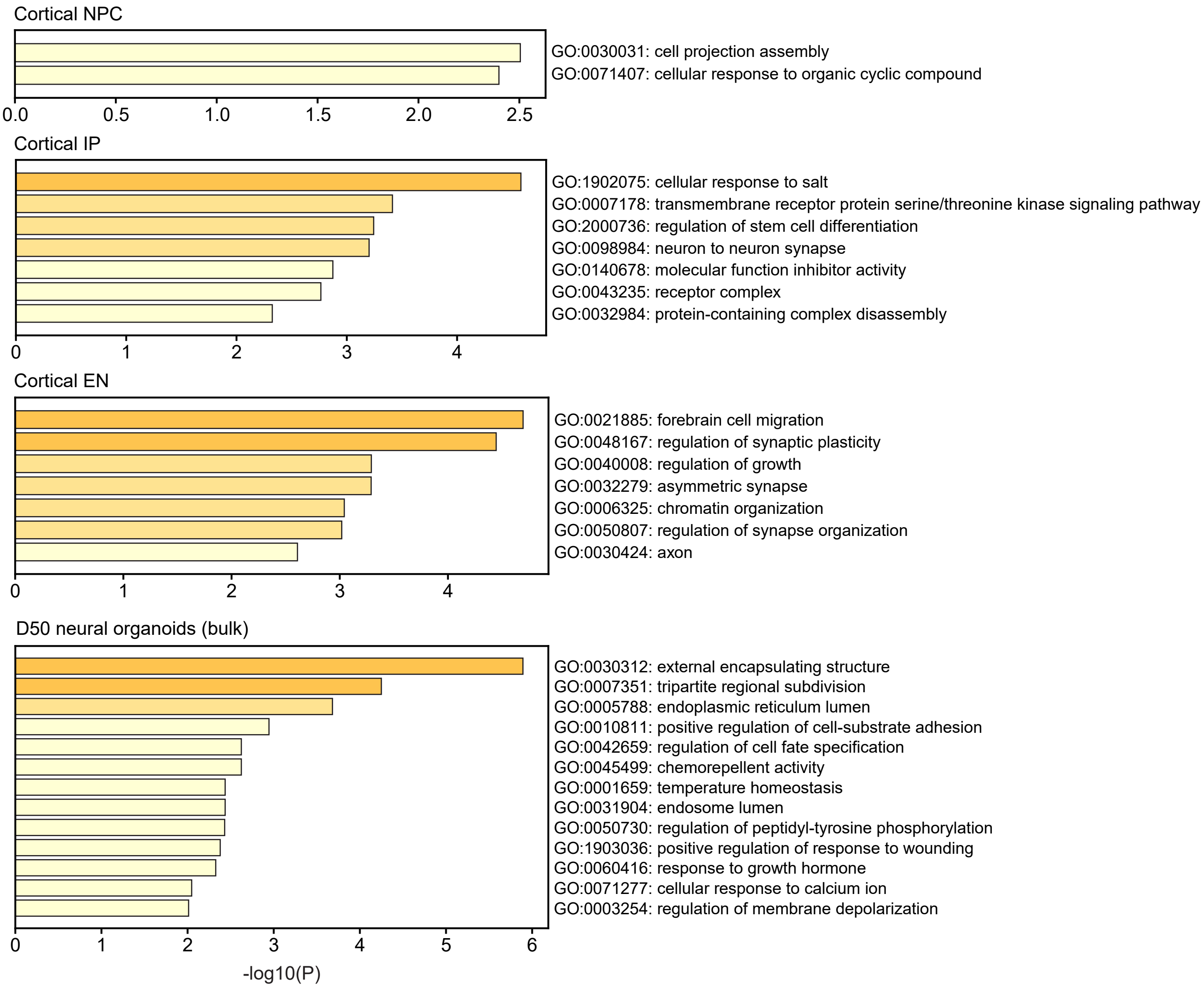


**Figure S14. Gene ontology enrichment analysis results for genes differentially expressed in 010A$^{KO/KO}$ compared to wild type neural organoids.** Bar graphs showing the results of GO enrichment analyses (biological processes, molecular functions and cellular processes) using Metascape for genes significiantly differentially expressed in the pseudo-bulk datasets of the dorsal cell types identified in the day-57 neural organoids and in the bulk transcriptomic data from day-50 organoids when the 010A$^{KO/KO}$ genotype was compared to wild type cells. All genes with a DESeq2 base mean expression value of > 0 were used as a background list.

**A**

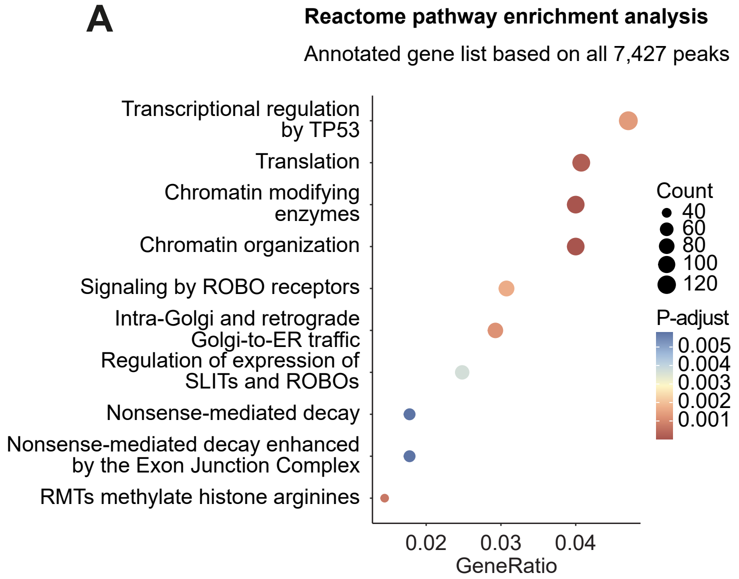


**B**

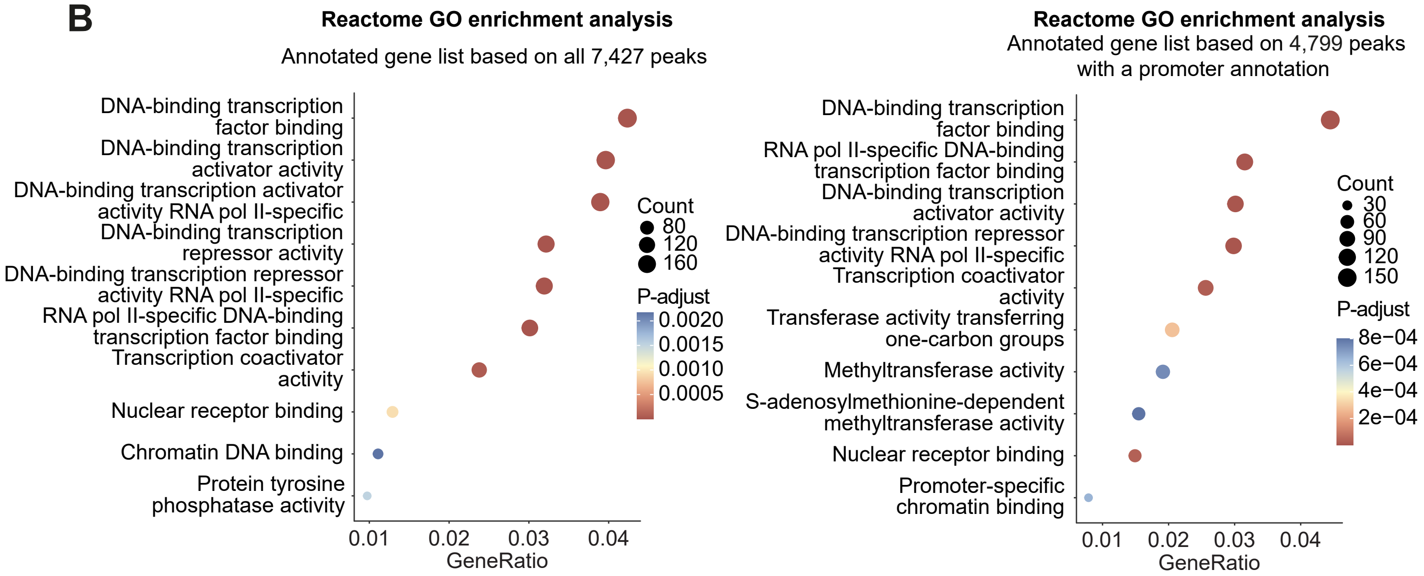


**C**

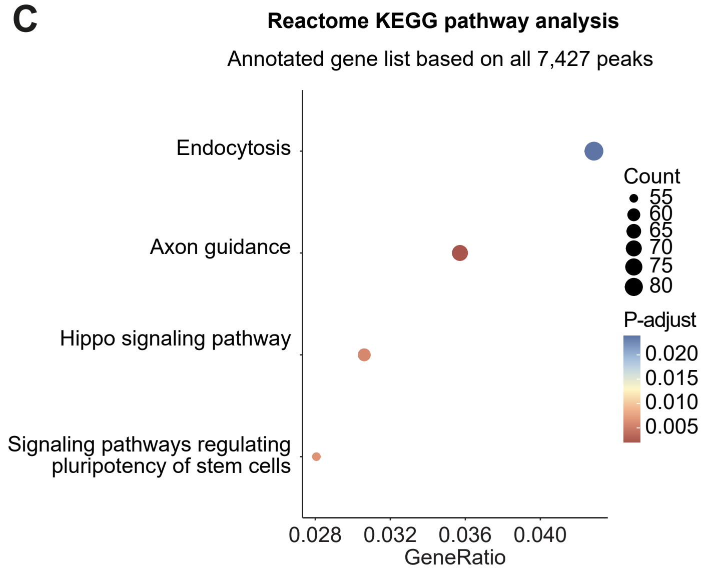


**Figure S15. Genes associated with CHD3 binding sites in unguided neural organoids are enriched for transcription regulation-related and axon guidance-related functions. A**) A dotplot showing the results of a Reactome pathway enrichment analysis of the genes associated with all high-confidence CHD3 ChIP-peaks. **B**) Dotplots showing the results of Reactome GO enrichment analyses. Left, enrichment for genes associated with all high-confidence CHD3 ChIP-peaks. Right, enrichment for genes associated with CHD3 ChIP-peaks with a promoter annotation only. **C**) A dotplot showing the results of a Reactome KEGG pathway enrichment analysis on the genes associated with all high-confidence CHD3 ChIP-peaks. For the genes associated with CHD3 ChIP-peaks with a promoter annotation only, no significant KEGG pathway enrichment was detected. **A-C**) The colouring from blue to red represents the adjusted *p* value, and the dot size the number of genes mapping to the terms.

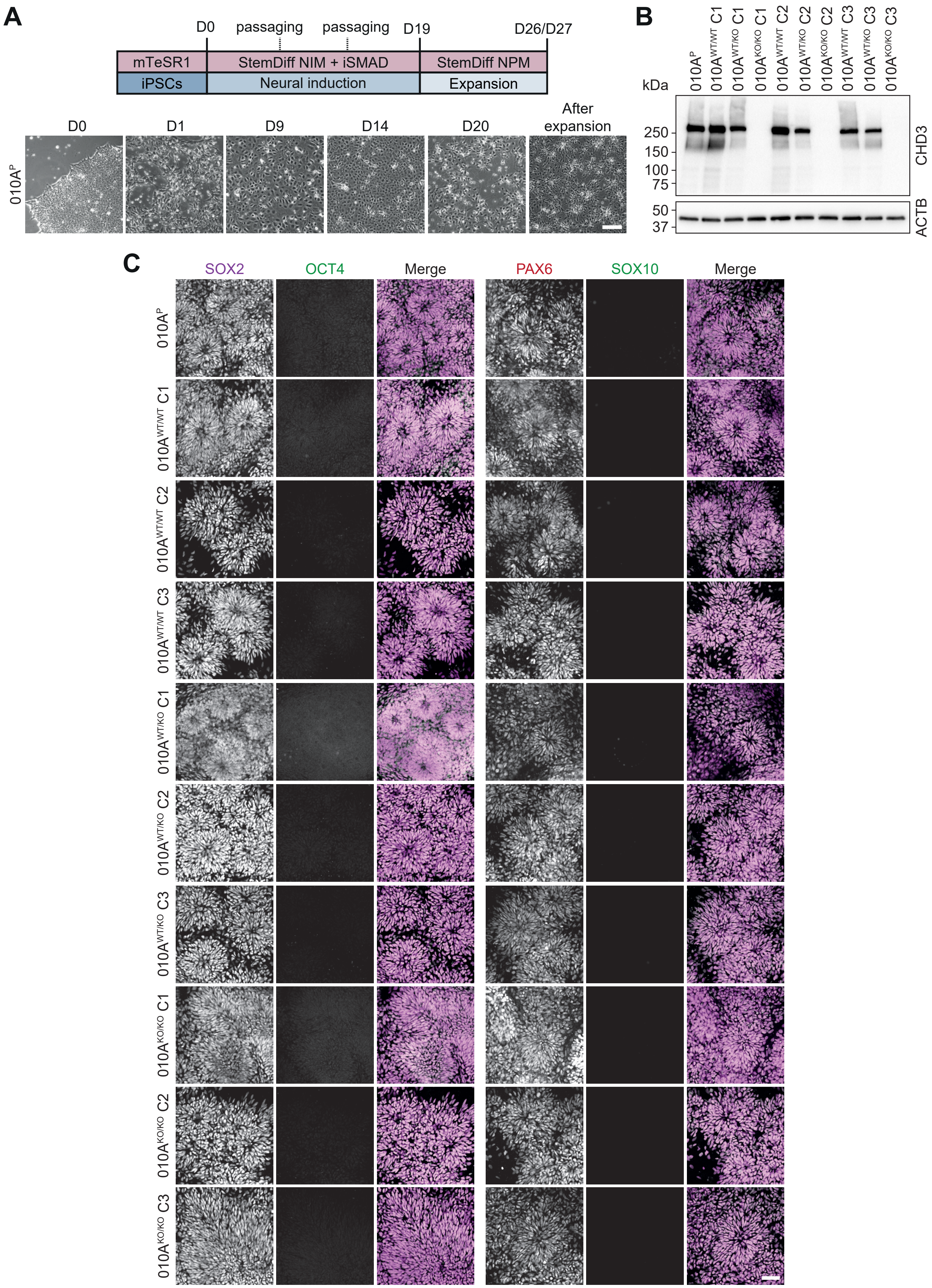


**Figure S16. Characterization of neural progenitor cell lines derived from the CHD3 CRISPR-Cas9 gene-edited iPSC lines. A**) Schematic of the protocol used to generate NPCs from iPSCs. Details are described in the Methods section. **B**) Immunoblot of whole-cell lysates of NPCs expressing CHD3 protein. Expected molecular weight is ~226 kDa. The blot was probed for ACTB to ensure equal protein loading. **C**) Immunohistochemistry micrographs of NPCs for the NPC markers SOX2 and PAX6 (magenta), pluripotency marker OCT4 (green) and neural crest cell marker SOX10 (green). Scale bar = 50 µm.

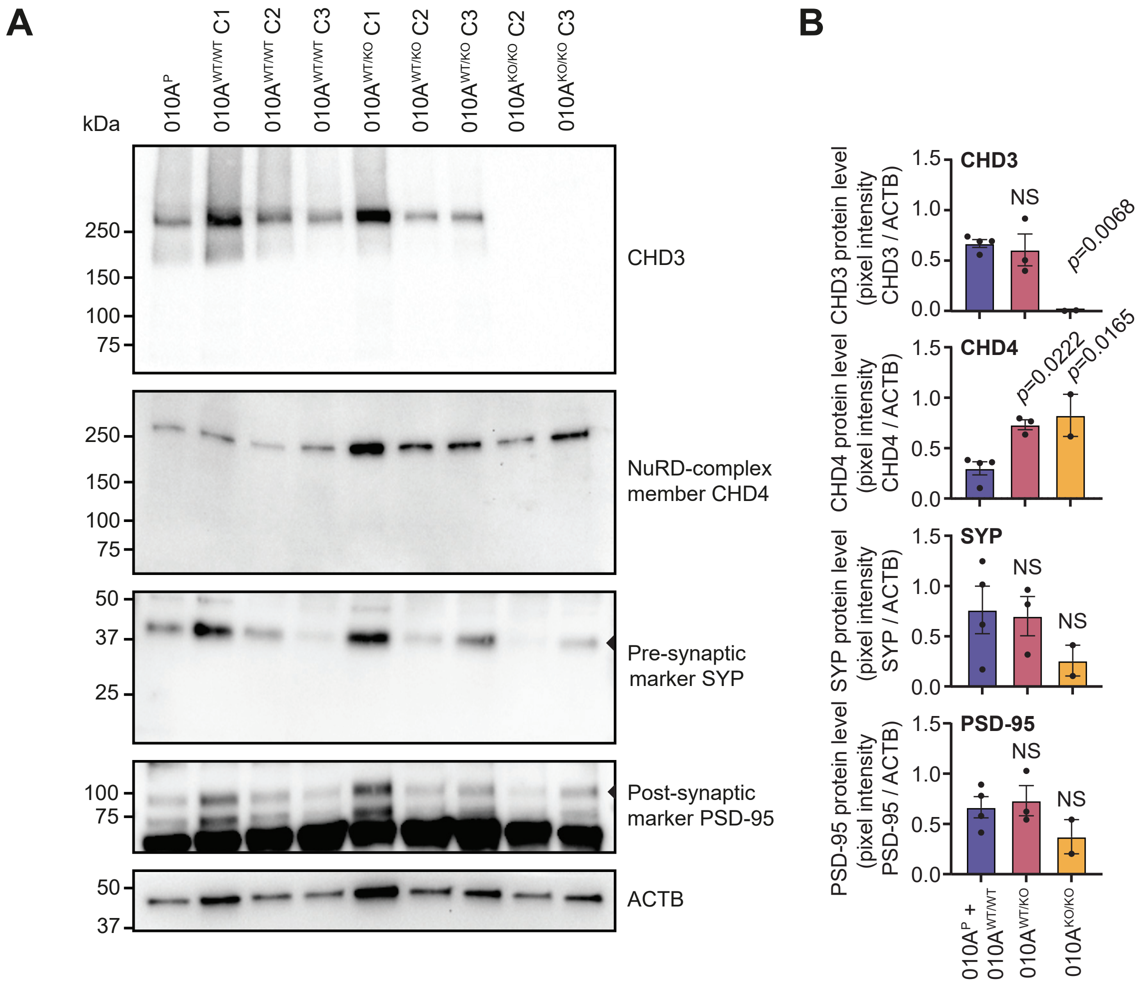


**Figure S17. Protein expression levels of proteins of interest in day-21 forebrain neurons derived from gene-edited iPSC lines.** **A**) Representative immunoblots of whole-cell lysates of D21 forebrain neurons. Expected molecular weights are CHD3: ~226 kDa, CHD4: ~218 kDa, SYP: ~38 kDa and PSD-95: ~85 kDa (although bands are typically detected at levels between 90-100 kDa due to phosphorylation). ACTB was used to ensure equal protein loading and to normalize for quantification of protein levels. **B**) Quantification of protein levels based on immunoblots shown in (**A**). For SYP and PSD-95, the bands used for quantification (at ~38 kDa and ~90-95 kDa respectively) are indicated with an arrow head in (**A**). For each protein of interest, two immunoblots were developed and the average pixel instensity values, normalized for the pixel intensity of ACTB, were calculated. The graphs show mean pixel intensity ratios across CHD3 genotypes with $n$ = 4 for wild-type samples (010A$^{P}$, 010A$^{WT/WT}$ C1, C2 and C3), $n$ = 3 for samples heterozygous for a CHD3 knockout allele (010A$^{WT/KO}$ C1, C2 and C3) and $n$ =2 for complete CHD3 knockouts (010A$^{KO/KO}$ C2 and C3). Conditions were compared to the wild-type samples using a one-way ANOVA followed by a *post hoc* Tukey test.

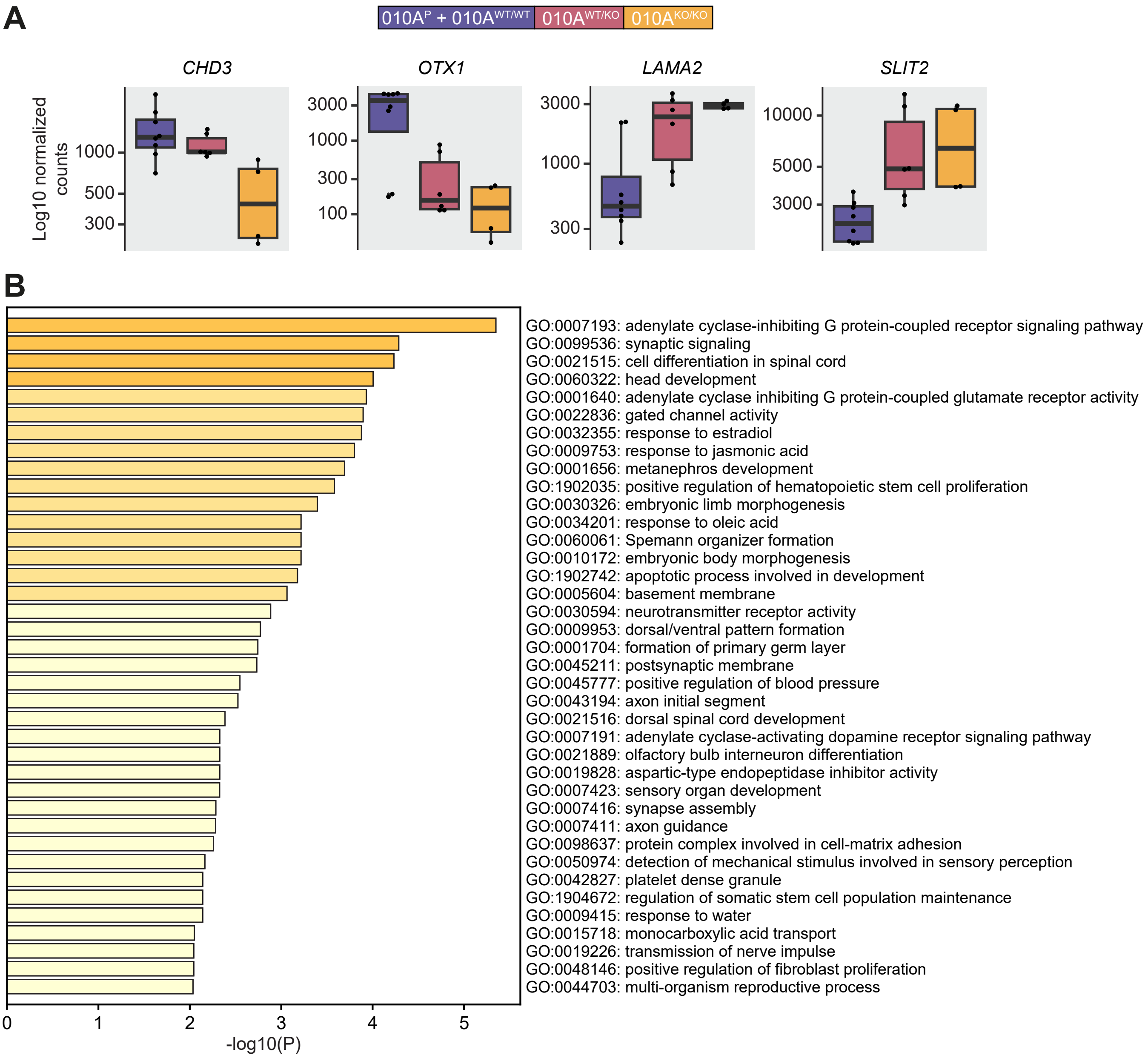


**Figure S18. The effects of *CHD3* disruption on gene expression in whole day-21 forebrain neurons based on bulk RNA sequencing data. A**) Box plots of the bulk transcriptomics normalized count data of whole day-21 forebrain neurons for a selection of differentially expressed genes (*CHD3*, *OTX1*, *LAMA2* and *SLIT2*) across *CHD3* genotypes. The 010A$^{P}$ and 010A$^{WT/WT}$ samples were shaded in purple, the 010A$^{WT/KO}$ samples in red, and the 010A$^{KO/KO}$ samples in yellow. **B**) A bar graph showing the results of a GO enrichment analysis (biological processes, molecular functions and cellular processes) using Metascape for genes significantly differentially expressed in day 21-forebrain neurons with a 010A$^{KO/KO}$ genotype compared to wild type cells. All genes with a DESeq2 base mean expression value of > 0 were used as a background list.

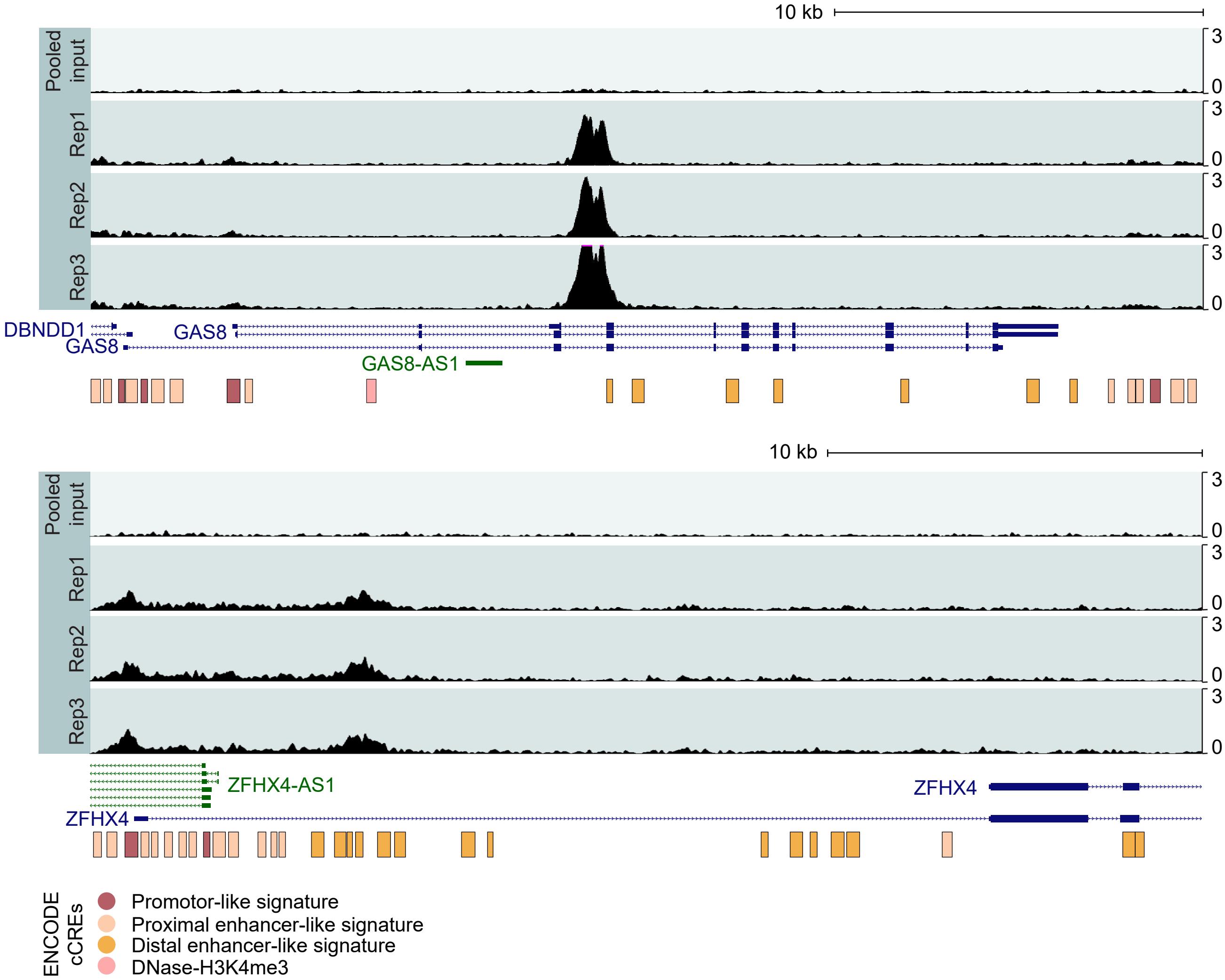


**Figure S19. CHD3 ChIP-seq peaks in the *GAS8* and *ZFHX4* genetic loci in day-50 neural organoids.** CHD3 enrichment across the genetic loci of *GAS8*/*GAS8-AS1* (top),and *ZFHX4*/*ZFHX4-AS1* (bottom) in day-50 neural organoids, with ENCODE Cis-Regulatory Elements (cCREs) annotated below. CHD3 ChIP-signals for the three independent replicates show high correlation, and the pooled input sample demonstrates absence of background in these regions.

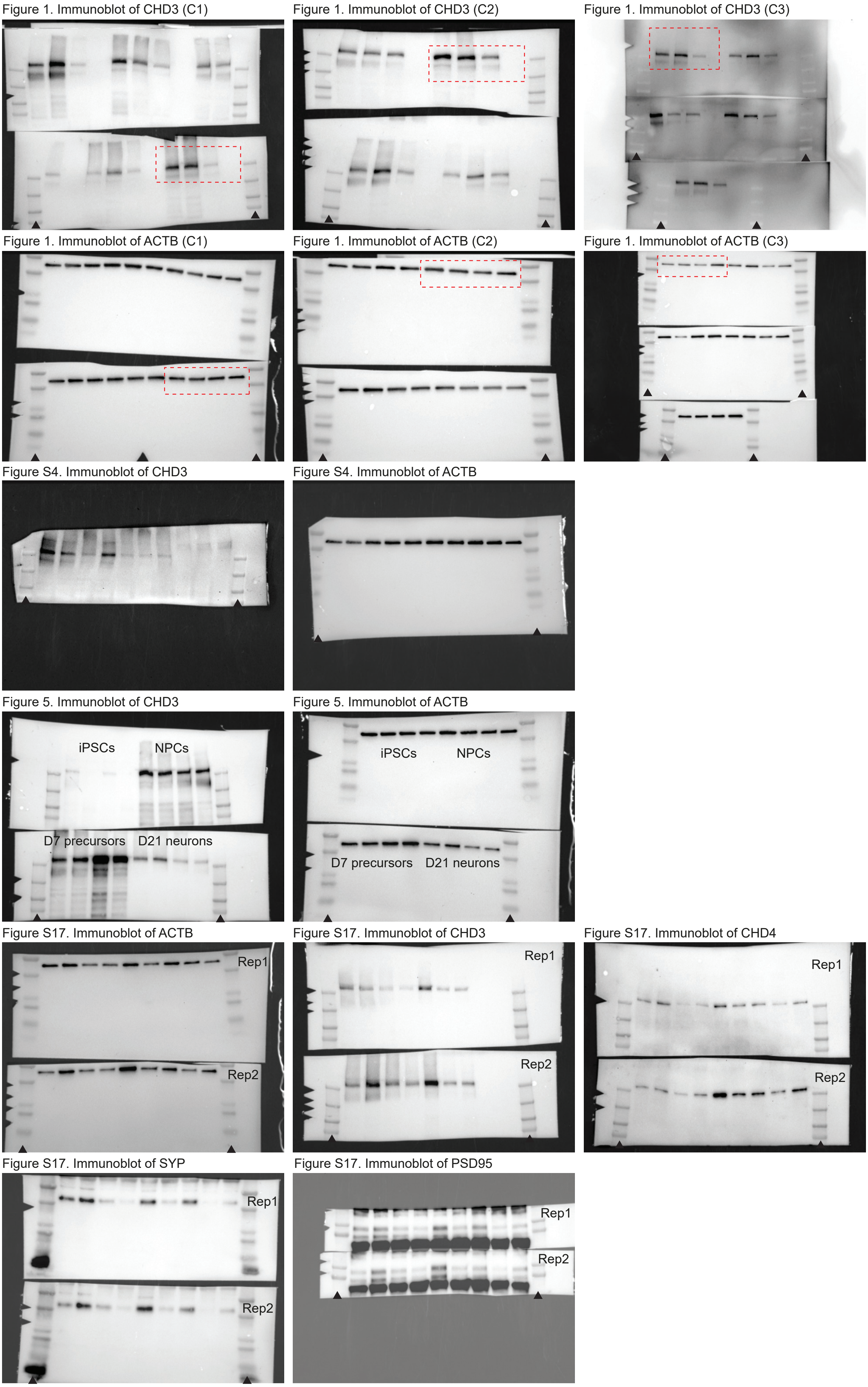


**Figure S20. Original uncropped images of immunoblots included in this study.** Arrow heads indicate the Precision Plus Protein Dual Color ladder (Bio-Rad). For immunoblots included in Figure 1, a dashed rectangle indicates which set of samples was selected for final presentation. For the other immunoblots, all samples were part of the final figures.
